# Brain transcriptome analysis reveals subtle effects on mitochondrial function and iron homeostasis of mutations in the *SORL1* gene implicated in early onset familial Alzheimer’s disease

**DOI:** 10.1101/2020.07.17.207787

**Authors:** Karissa Barthelson, Stephen Martin Pederson, Morgan Newman, Michael Lardelli

## Abstract

**Background:** To prevent or delay the onset of Alzheimer’s disease (AD), we must understand its molecular basis. The great majority of AD cases arise sporadically with a late onset after 65 years of age (LOAD). However, rare familial cases of AD can occur due to dominant mutations in a small number of genes that cause an early onset prior to 65 years of age (EOfAD). As EOfAD and LOAD share similar pathologies and disease progression, analysis of EOfAD genetic models may give insight into both subtypes of AD. Sortilin-related receptor 1 (*SORL1*) is genetically associated with both EOfAD and LOAD and provides a unique opportunity to investigate the relationships between both forms of AD. Currently, the role of *SORL1* mutations in AD pathogenesis is unclear.

**Methods:** To understand the molecular consequences of *SORL1* mutation, we performed targeted mutagenesis of the orthologous gene in zebrafish. We generated an EOfAD-like mutation, V1482Afs, and a putatively null mutation, to investigate whether EOfAD-like mutations in *sorl1* display haploinsufficiency by acting through loss-of-function mechanisms. We performed mRNA-sequencing on whole brains comparing normal (wild type) fish with their siblings heterozygous for EOfAD-like or complete loss-of-function mutations in *sorl1* or transheterozygous for these mutations. Differential gene expression and gene set enrichment analyses identified, respectively, changes in young adult zebrafish brain transcriptomes, and putative effects on neural subcellular functions.

**Results:** We identified subtle effects on expression of genes involved in energy production, mRNA translation and mTORC1 signalling in both the EOfAD-like and null mutant brains, implying that these effects are due to *sorl1* haploinsufficiency. Surprisingly, we also observed changes to expression of genes occurring only in the EOfAD-mutation carrier brains, suggesting gain-of-function effects. Transheterozygosity for the EOfAD-like and null mutations (i.e. lacking wild type *sorl1*), caused apparent effects on iron homeostasis and other transcriptome changes distinct from the single-mutation heterozygous fish.

**Conclusions:** Our results provide insight into the possible early brain molecular effects of an EOfAD mutation in human *SORL1*. Differential effects of heterozygosity and complete loss of normal *SORL1* expression are revealed.

## Background

Alzheimer’s disease (AD) is a progressive neurodegenerative disorder and the most common form of dementia. AD brains show a broad range of pathologies including deposition of intercellular deposits of insoluble amyloid β (Aβ) peptides within plaques, intracellular tangles primarily consisting of hyperphosphorylated tau proteins, vascular abnormalities (1, 2), mitochondrial dysfunction (3–5), inflammation (6, 7), lipid dyshomeostasis (8–10), metal ion dyshomeostasis (11, 12) and numerous others.

To prevent or delay the onset of AD, we need to understand the early cellular changes which eventually lead to these AD pathologies. This is difficult to investigate in humans, as pre-symptomatic, living AD brain tissue is inaccessible for detailed molecular analysis. Consequently, animal models can be extremely useful for understanding the stresses driving this disease. The commonly used mouse models of AD overexpress human mutant forms of the EOfAD genes to show histopathological phenotypes reminiscent of the human disease. Troublingly however, the brain transcriptomes of these mouse models show low concordance with human AD, and with each other (13). Thus, these mouse models are unlikely to mimic, accurately, the genetic state of the human disease.

In rare, familial cases of AD, patients can show symptoms of disease onset before 65 years of age. These cases are most often due to single, autosomal dominant mutations in one of three genes: amyloid β A4 precursor protein (*APP*), presenilin 1 (*PSEN1*) and presenilin 2 (*PSEN2*) (EOfAD) (14). All EOfAD mutations in the *PSENs* and *APP* follow a “reading-frame preservation rule”, where mutations causing truncation of the open reading frame do not cause EOfAD (reviewed in (15)). Recently, mutations in sortilin-related receptor 1 (*SORL1*) have been found that segregate with early onset AD in dominant inheritance patterns (16, 17), suggesting that *SORL1* may represent a fourth EOfAD-causative gene. Interestingly, both missense and reading frame-truncating mutations in *SORL1* have been observed in early-onset AD families (16, 17). Since reading frame-truncating mutations in *SORL1* have been shown likely to cause nonsense mediated decay of the mutant alleles’ mRNA (16), it is thought the mutations may act through a haploinsufficient, loss-of-function mechanism. However, this has not been explored at the molecular level *in vivo*.

Most AD cases arise sporadically and have an age of onset of later than 65 years (LOAD). The etiology of LOAD is still unclear. However, there are variants at numerous loci associated with increased risk of developing LOAD, such as the ε4 allele of the gene encoding apolipoprotein E (*APOE*) (18). Interestingly, variation at the *SORL1* locus is also associated with LOAD (19–22) so that understanding the function of this gene may illuminate a mechanistic link between EOfAD and LOAD. However, whether or not *SORL1* should be regarded as an EOfAD-causative locus in the manner of the *PSEN* and *APP* genes is still debated (23).

SORL1 protein is a membrane-bound, multi-domain-containing protein and localises mainly in cells’ endolysosomal system and the trans-Golgi network. We have previously found that SORL1 localises to the mitochondrial associated membranes (MAMs) of the endoplasmic reticulum (24). SORL1 belongs to the family of vacuolar protein sorting 10 (VPS10)-containing proteins, or sortilins. These proteins all carry a VPS10 domain with homology to the VPS10P domain found in yeast (25). SORL1 also belongs to the low density lipoprotein receptor (LDLR) family of proteins and contains both LDLR class A repeats and LDLR class B repeats (reviewed in (26)).

The functional role of SORL1 has been investigated mostly in the context of the distribution and processing of APP within cells. SORL1 guides APP throughout the endolysosomal system and is thought to prevent formation of Aβ by promoting recycling of APP (reviewed in (26)). Mutations in the protein-coding region of *SORL1* have been shown to reduce the capacity of the SORL1 protein to bind APP and result in increased levels of Aβ (27, 28). A single nucleotide polymorphism (SNP) cluster consisting of 6 SNPs spanning a region between exon 6 to intron 9 of *SORL1* is associated with decreased elevation of *SORL1* expression in response to brain-derived neurotrophic factor (BDNF) in neurons derived from human induced pluripotent stem cells (hiPSCs). This results in aberrant processing of APP (29)). However, the non-APP related functions of *SORL1,* and the effects of mutations in *SORL1* on the molecular state of the central nervous system *in vivo,* remain largely unexplored.

Here, we describe a study addressing two questions: 1) What are the effects of heterozygosity for an EOfAD-like mutation in *sorl1* on the transcriptomes of young-adult, mutation carrier brains *in vivo*? 2) Are the effects of a heterozygous EOfAD-like mutation in *sorl1* due to a loss- or gain-of-function? To address these questions, we introduced an EOfAD-like mutation, V1482Afs, into the zebrafish orthologue of *SORL1* (*sorl1*) to mimic the human EOfAD mutation C1478* (16). We also generated a putatively null mutation, R122Pfs, as a control representing haploinsufficient loss-of-function. We performed RNA sequencing (RNA-seq) on mRNAs derived from entire brains from a family of young-adult sibling zebrafish that were either heterozygous for the V1482Afs mutation (hereafter referred to as EOfAD-like/+), heterozygous for the null mutation (hereafter referred to as null/+), transheterozygous for both the null and EOfAD-like mutations (*i.e.* EOfAD/null, a complete loss of wild type *sorl1*) or wild type. We found that, in the heterozygous state, the EOfAD-like mutation causes subtle changes to gene expression and appears to act through both loss-of-function and gain-of-function mechanisms. Differences in the EOfAD-like/+, null/+ and transheterozygous mutant sibling brain transcriptomes highlight the importance of analysing animal models which reflect, as closely as possible, the genetic state of the human disease, and illuminate novel cellular processes previously unknown to be affected by loss of normal *SORL1* expression.

## Methods

### Zebrafish husbandry and animal ethics

Work with zebrafish was performed under the auspices of the Animal Ethics Committee of the University of Adelaide, permit numbers S-2017-089 and S-2017-073. All zebrafish used in this study were maintained in a recirculating water system on a 14 hour light/10 hour dark cycle, and fed NRD 5/8 dry food (Inve Aquaculture, Dendermonde, Belgium) in the morning and live *Artemia salina* in the afternoon. In total, 36 fish were used over all the experiments described in this study.

### Genome editing constructs

To introduce an EOfAD-like frameshift mutation near the C1481 codon of zebrafish *sorl1*, we used a TALEN pair designed by, and purchased from, Zgenebio Biotech Inc. (Taipei City, Taiwan). The genomic DNA recognition sites (5’ to 3’) were TGAGGTGGCGGTGTG (left TALEN) and CTGAAATACATGCTGG (right TALEN) (**Additional File 1**). The DNAs encoding the TALEN pair protein sequences were supplied in the pZGB2 vector. These constructs were linearised with *Not* I (NEB, Ipswich, USA) and then mRNA was transcribed *in vitro* using the mMESSAGE mMACHINE T7 *in vitro* transcription kit (Invitrogen, Carlsbad, USA) following the manufacturer’s protocol.

To generate a putatively null mutation in *sorl1*, we targeted exon 2 of *sorl1* using the CRISPR-Cpf1 system (30). A crRNA was designed to recognise the sequence 5’ GGGGTCTGTGGCACCAACGCT 3’ in exon 2 of *sorl1,* with a PAM sequence of TTTT (**Additional File 1**). Both the crRNA and Cpf1 recombinant protein were synthesised by, and purchased from, IDT Technologies (Iowa, USA).

We injected these constructs into zebrafish embryos at the one-cell stage and, ultimately, isolated fish carrying the mutations of interest (V1482AfsTer12, hereafter referred to as V1482Afs or “EOfAD-like”, in exon 32 and R122PfsTer118, hereafter referred to as R122Pfs or “null”, in exon 2). For a detailed description of the methods used to isolate these mutant lines of fish, see **Additional File 2.**

### Breeding strategy

To generate families of fish for analysis, we crossed an EOfAD-like/+ fish to a null/+ fish to generate a family of siblings with one of four *sorl1* genotypes: EOfAD-like/+, null/+, EOfAD-like/null (transheterozygous) or +/+ (**Figure 1**). Each family of fish was raised in an 8 litre capacity tank, with approximately 40 fish per tank (i.e. approximately 5 fish per litre) to avoid stresses due to overcrowding. The tanks were placed side-by-side in the same recirculated water aquarium system to reduce environmental variation between them.

**Figure 1:**
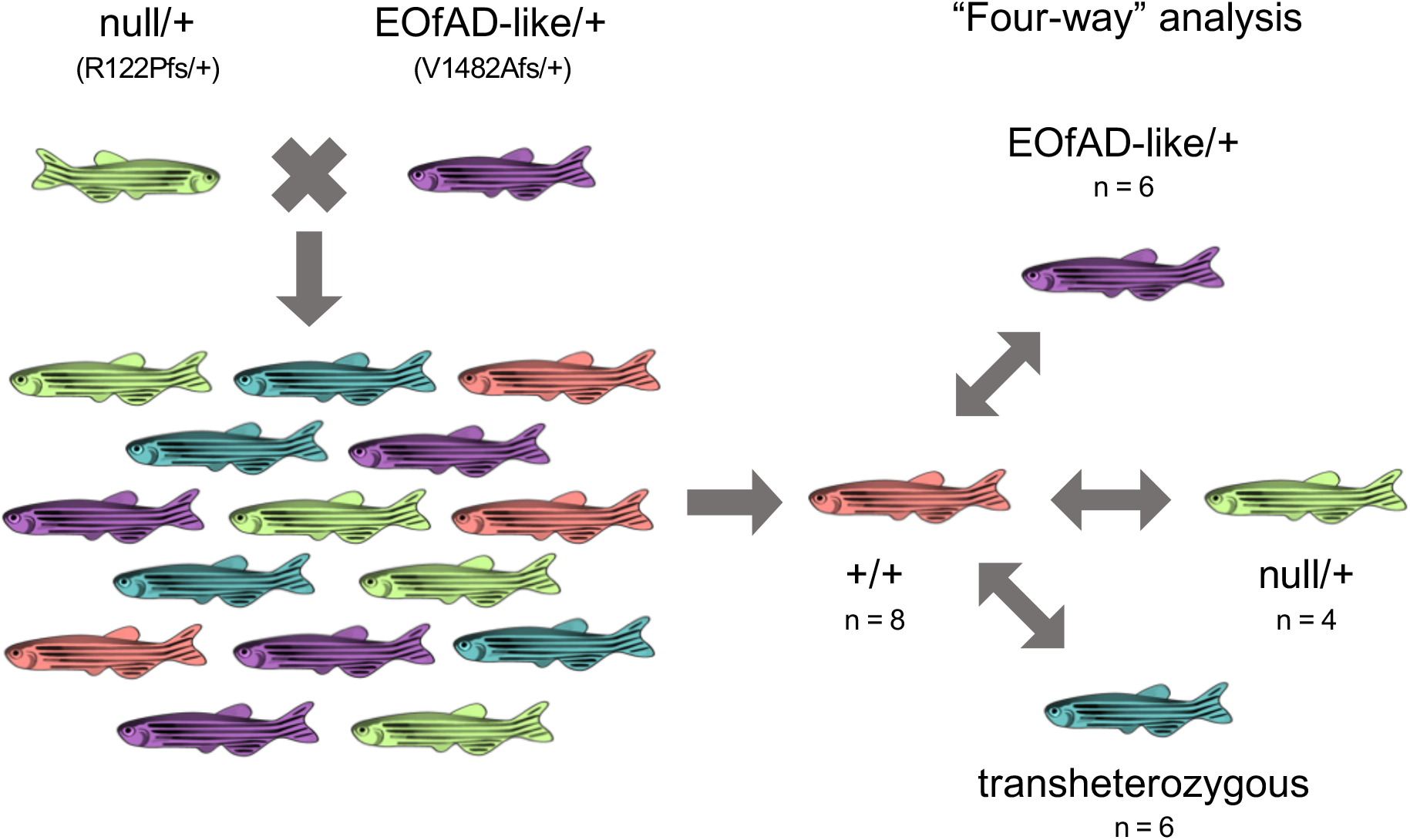
Breeding strategy We mated a pair of fish with *sorl1* genotypes null/+ and EOfAD-like/+ to generate families of fish composed of *sorl1* genotypes EOfAD-like/+, null/+, EOfAD-like/null (transheterozygous) or wild type (+/+). The values of n are representative of the numbers of fish used in the RNA sequencing experiment.

### Allele-specific expression of *sorl1* transcripts

Adult zebrafish (6 months old) were euthanised in a loose ice slurry. Entire heads were cut off and placed in 600 μL of RNAlater™ Stabilization Solution (Invitrogen, Carlsbad, USA) before incubation at 4 °C overnight. Brains were then removed from the heads using sterile watchmaker’s forceps. Total RNA was extracted from the brains using the QIAGEN RNeasy® Mini Kit (Qiagen, Venlo, Netherlands) according to the manufacturer’s protocol. Recovered RNA concentrations were estimated using a Nanodrop spectrophotometer. RNAs were stored at −80 °C until required. We prepared cDNA using 400ng of total RNA input (to give a 20 ng/μL cDNA solution) and the Superscript III First Strand Synthesis System (Invitrogen, Carlsbad, USA) according to the manufacturer’s instructions (random hexamer priming method).

We quantified the absolute copy numbers of *sorl1* transcripts in EOfAD-like/+ mutant brains by allele-specific digital quantitative PCRs (dqPCRs) as described in (31). The mutant allele-specific forward primers described in **Additional File 2** were used with a common reverse primer binding over the *sorl1* exon 33-34 junction (5’ GACCTGCTGTTCATCAGTGC 3’) to avoid amplification from any genomic DNA carried over during the RNA extraction. We used 50 ng of cDNA as input in each dqPCR reaction.

To compare the *sorl1* transcript expression levels in null/+ mutant brains, we amplified a ~250 bp region spanning the R122 codon in exon 2 from 50 ng of brain-derived cDNA from wild type and null/+ fish using reverse transcription PCRs (RT-PCRs) and resolved the RT-PCR products on a 3% agarose gel in 1 x tris-acetate-EDTA (TAE) in milliQ. Since the null mutation deletes 16 bp from the gene, this procedure resolved the two bands representing the two alleles of *sorl1* and we could inspect the relative intensity of the bands.

### RNA sequencing data generation and analysis

We performed RNA sequencing in the “four-way” analysis described in **Figure 1.** Adult zebrafish (6 months old) were euthanised in an ice slurry. Whole heads were removed for preservation in RNAlater solution (Invitrogen, Carlsbad, USA), and a fin biopsy was removed for genotype determination by PCRs (**Additional File 2**). The whole brains were removed from the preserved heads using sterile watchmaker’s forceps and then total RNA was isolated using the mirVana™ miRNA Isolation Kit (Ambion, Life Technologies, Thermo Fisher Scientific, Waltham, USA) following the manufacturer’s protocol. Genomic DNA was removed from the total RNA by DNase treatment using the DNA-free™ Kit (Ambion, Life Technologies, Thermo Fisher Scientific, Waltham, USA).

500 ng of total RNA was then delivered to the Genomics service at the South Australian Health and Medical Research Institute (SAHMRI, Adelaide, AUS) for poly-A+, stranded library preparation and RNA sequencing using the Illumina Nextseq platform. We used a total of 24 fish at 6 months of age (EOfAD-like/+ : n = 6, null/+: n = 4, wild type: n = 8 and transheterozygous: n = 6). We initially aimed to have n = 6 fish per genotype, based on a previous power calculation indicating that n = 6 would provide approximately 70% power to detect the majority of expressed transcripts in a zebrafish brain transcriptome at a fold-change > 2 and at a false discovery rate of 0.05 (data not shown). However, genotype checks of the RNA-seq data revealed that two samples had been misidentified during the genotyping by PCRs.

Demultiplexed fastq files were provided by SAHMRI as 75bp single-end reads. All libraries were sequenced to a depth of between 30 and 38 million reads, across four NextSeq lanes which were subsequently merged. The quality of the provided reads was checked using fastQC (32) and ngsReports (33). We then trimmed adaptors and bases with a PHRED score of less than 20 from the reads using AdapterRemoval (v2.2.0). Reads shorter than 35 bp after trimming were discarded. We then performed a pseudo-alignment to estimate transcript abundances using kallisto v0.43.1 (34) in single-end mode, specifying a forward stranded library, the fragment length as 300 with a standard deviation as 60, and 50 bootstraps. The index file used for the kallisto pseudo-alignment was generated from the zebrafish transcriptome according to the primary assembly of the GRCz11 reference (Ensembl release 96), with the sequences for unspliced transcripts additionally included.

We imported the transcript abundances from kallisto for analysis using R (35) using the catchKallisto function from the package edgeR (36). We summed the counts of each of the mature transcripts arising from a single gene (i.e. omitting any unspliced transcripts with intronic sequences remaining) to generate gene-level quantifications of transcript abundances. We filtered genes which were undetectable (less than 0.66 counts per million reads (CPM)) in at least 12 of the 24 RNA-seq libraries, leaving library sizes ranging between 13,481,192 and 22,737,049 counts.

Principal component analysis identified that the largest source of variation in the RNA-seq dataset was due to library size (discussed later). Therefore, to assist in identification of changes to gene expression due to *sorl1* genotype, we corrected for this unwanted variation using the RUVg method of RUVSeq (37). To identify the negative control genes for RUVg, we performed an initial differential gene expression analysis using the generalised linear model capabilities of edgeR in an ANOVA-type method. A design matrix was specified with the wild type *sorl1* genotype as the intercept, and the coefficients as each of the other *sorl1* genotypes (EOfAD-like/+, null/+, EOfAD-like/null), the sex of the fish (male or female, with female as the reference level) the tank in which each fish was raised (tank 1, tank 2 or tank 3, with tank 1 being the reference level). The 5000 least differentially expressed genes (i.e. the genes with the highest p-value) due to all *sorl1* genotypes were then used as the negative control genes to remove one factor of unwanted variation using RUVSeq. The resulting W_1 offset term, setting *k* = 1, from RUVSeq was included in the design matrix for an additional differential gene expression analysis. We considered a gene to be differentially expressed (DE) if the FDR-adjusted p-value was below 0.05 for each specific comparison.

We also checked within the initial differential expression analysis to determine whether there was a bias for GC content or length with the fold change or significance of each gene. No discernible length or GC bias was found (**Additional File 3**).

For enrichment analysis, the gene sets used were the HALLMARK (38) and KEGG pathway (39) gene sets available from the Molecular Signatures Database (MSigDB, www.gsea-msigdb.org/gsea/msigdb/index.jsp). We downloaded these gene sets as a .gmt file with human Entrez gene identifiers and converted the Entrez identifiers to zebrafish Ensembl identifiers using a mapping file obtained from the Ensembl biomart (40) web interface. We also used the four gene sets of genes with an iron-response element (IRE) described in (41) to determine whether there was a possible iron dyshomeostasis signal in our dataset. Finally, the GROSS_HYPOXIA_VIA_HIF1A_DN and GROSS_HYPOXIA_VIA_HIF1A_UP (42) gene sets from MSigDB (C2, CPG subcategory) were used to characterise any changes to expression of genes involved in the cellular response to hypoxia.

Enrichment testing was performed using fry (43) and camera (44) from the limma package (45), and the fast implementation of the GSEA algorithm described in (46) using fgsea (47) for each comparison between the *sorl1* mutants and their wild type siblings.

Since each algorithm of enrichment analysis gave different levels of significance for each gene set, we calculated a consensus p-value by calculating their harmonic mean p-value (48). Gene sets were considered to be significantly altered as a group if the FDR-adjusted harmonic mean p-value was less than 0.05.

Visualisation for the RNA-seq analysis was performed using ggplot2 (49), pheatmap (50) and upsetR (51).

## Results

### Creation of *in vivo* animal models of null and EOfAD-like mutations in *SORL1*

The C1478* mutation in *SORL1* was identified in a French family and appears to segregate with AD with an autosomal dominant inheritance pattern (16). We aimed to generate a zebrafish model of this mutation by editing the endogenous sequence of zebrafish *sorl1*. The C1478 codon is conserved in zebrafish, C1481 (**Figure 2**). Consequently, we used TALENs to generate double stranded breaks at this site in the zebrafish genome (**Additional File 1**) and then allowed the non-homologous end-joining pathway of DNA repair to generate indel mutations. We identified a family of fish carrying a 5 nucleotide deletion, causing a frameshift in the coding sequence. This results in 11 novel codons followed by a premature termination codon (V1482AfsTer12, or more simply, V1482Afs, hereafter referred to as “EOfAD-like”).

**Figure 2:**
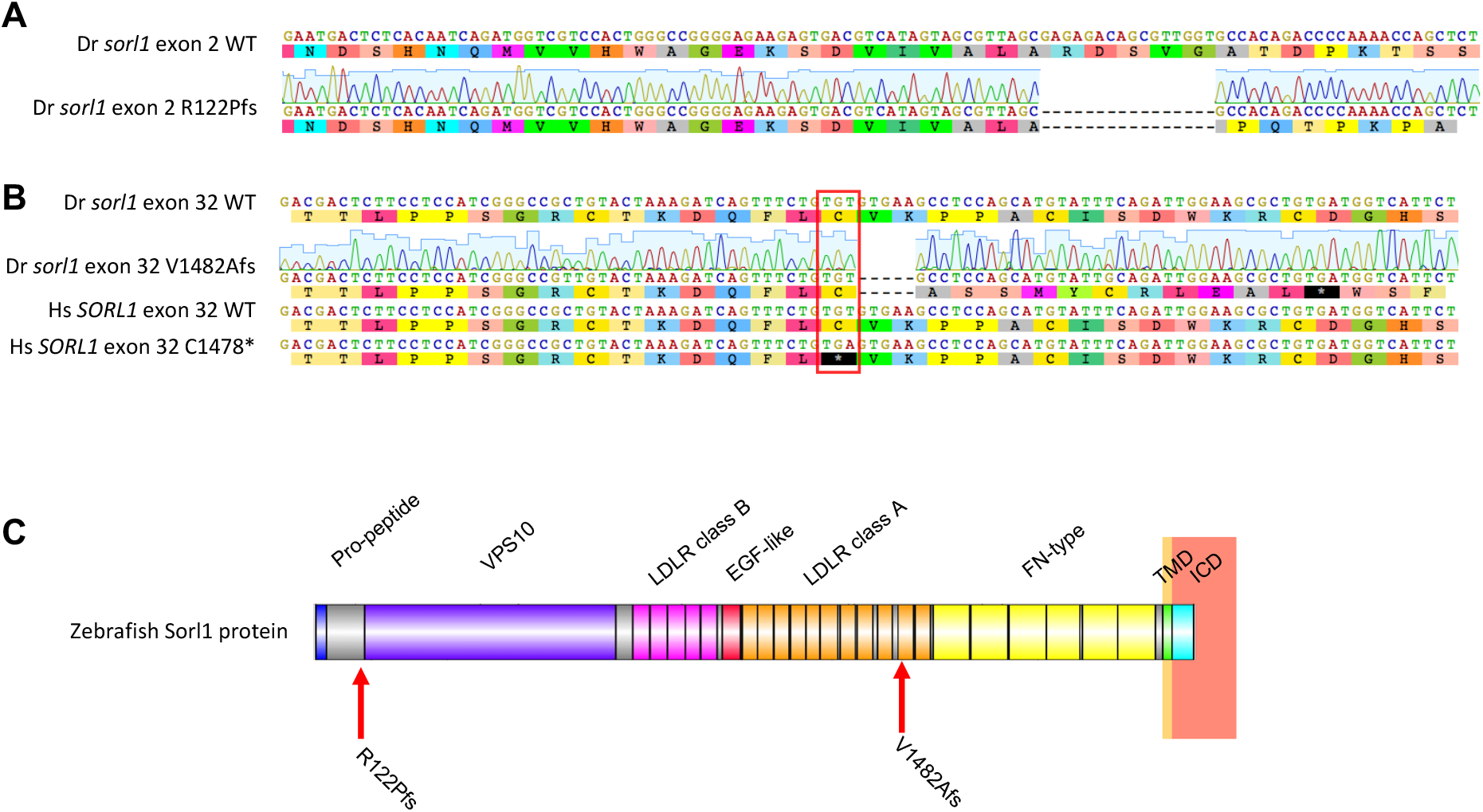
Mutations generated in zebrafish *sorl1* **A.** Zebrafish (Dr) *sorl1* wild type (WT) and null (R122Pfs) exon 2 sequences with the chromatogram showing the Sanger sequencing of the null allele. Both the DNA and amino acid sequences are shown. **B**. Alignment of the WT and EOfAD-like (V1482Afs) zebrafish *sorl1* exon 32 sequences, with the chromatogram showing Sanger sequencing of the EOfAD-like allele of *sorl1*. The human (Hs) *SORL1* exon 32 region is also shown with the C1478* mutation site and the equivalent C1481 zebrafish codon highlighted in red. **C**. A schematic of the zebrafish Sorl1 protein with the protein domains and mutation sites indicated. VPS10: vacuolar protein sorting 10 domain. LDLR: low density lipoprotein receptor. EGF: epidermal growth factor. FN: fibronectin. TMD: transmembrane domain. ICD: intracellular domain. Hs denotes *Homo sapiens*. Dr denotes *Danio rerio*.

To investigate whether the EOfAD-like mutation acts through loss-of-function, we generated a putatively null mutation in *sorl1*. We targeted exon 2 of *sorl1* using the CRISPR-Cpf1 system, as exon 1 contained three in-frame ATG codons which could, potentially, act as alternative translation initiation codons to allow translation of the majority of the protein. We identified a family of fish carrying a 16 nucleotide deletion, resulting in a frame shift in the coding sequence. This frame shift is predicted to encode 118 novel amino acids followed by a premature termination codon (R122PfsTer118, or more simply, R122Pfs, hereafter referred to as “null”). The protein encoded by the null allele of *sorl1* is predicted to lack most of the functional domains of the wild type Sorl1 protein (**Figure 2**).

### Protein-truncating mutations in *sorl1* are likely subject to nonsense-mediated mRNA decay

The C1478* mutation in human *SORL1* has been shown likely to cause nonsense mediated mRNA decay (NMD) in lymphoblasts from a human mutation-carrier (16). Therefore, we investigated whether transcripts of the EOfAD-like allele of *sorl1* are also subject to NMD by allele-specific digital quantitative PCRs (dqPCRs) on cDNA generated from EOfAD-like/+ brains. We observed significantly fewer transcripts of the EOfAD-like allele than the wild type allele in EOfAD-like/+ brains (p = 0.05). We also observed significantly fewer (p = 0.001) transcripts of the wild type allele in the EOfAD-like/+ brains than in their wild type siblings (**Figure 3A**).

**Figure 3:**
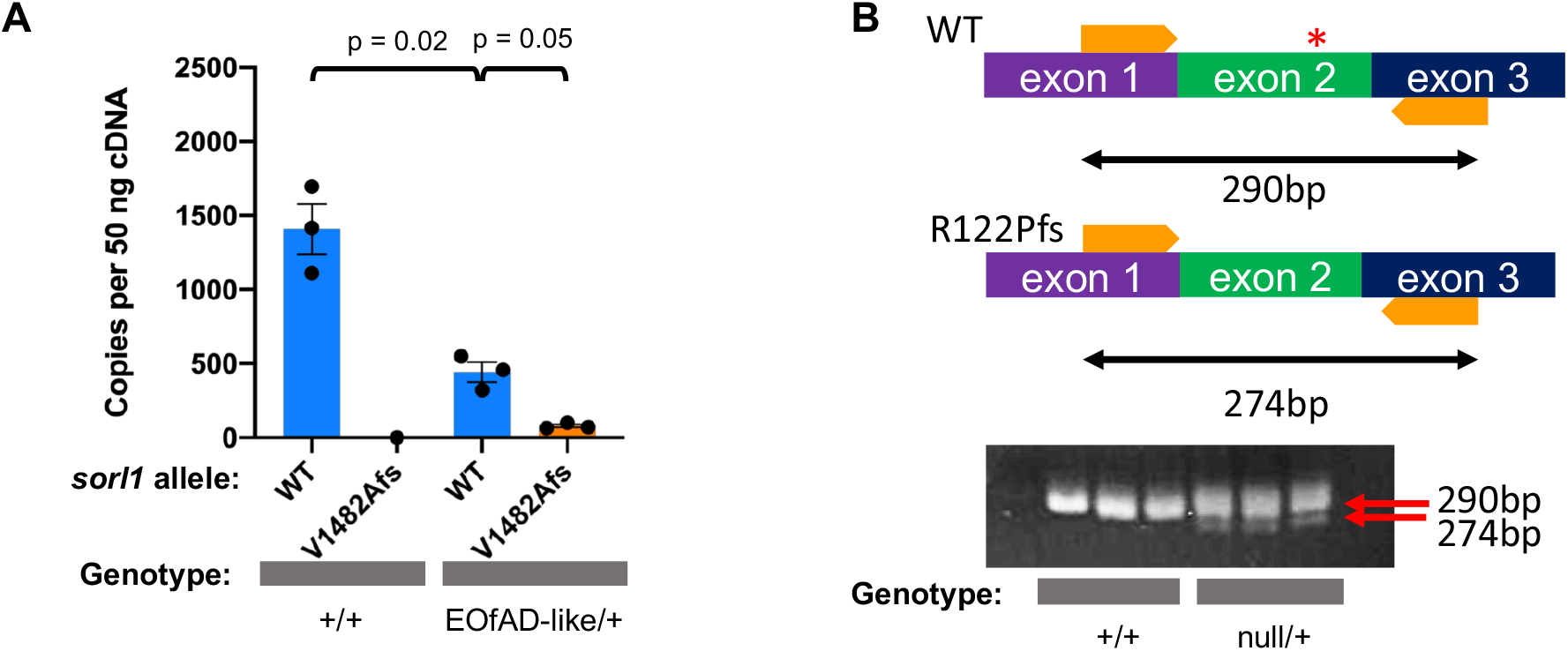
Allele-specific expression of *sorl1* transcripts **A**. The number of copies of *sorl1* transcripts present in +/+ (n = 3) and EOfAD/+ mutant (n = 3) brains per 50 ng of brain-derived cDNA. Data is presented as the mean ± SEM. In the +/+ brains, there were approximately 1400 copies of the wild type (WT) *sorl1* transcript and no copies of the EOfAD-like (V1482Afs) allele transcript. In the EOfAD-like/+ brains, there were approximately 440 copies of the WT *sorl1* transcript, and approximately 80 copies of the EOfAD-like allele transcript. P-values were determined by a one-way ANOVA with Dunnett’s T3 post-hoc test. **B.** A schematic of *sorl1* mRNA from exons 1 to 3. Primer binding sites are indicated by orange arrows, and the mutation site by the red star. Below is an image captured after agarose gel electrophoresis of the RT-PCR products. Lanes 1-3 are amplifications from three +/+ brains, while lanes 4-6 are amplifications from three null/+ brains. All fish were siblings.

We were also interested to determine whether the null transcript of *sorl1* was subject to NMD. It proved difficult to obtain differential amplification distinguishing the null and wild type alleles of *sorl1* in dqPCRs. Therefore, we used reverse transcription PCRs (RT-PCRs) to amplify a 290 bp region encompassing *sorl1* exon 2 from 50 ng of brain-derived cDNAs from null/+ fish and +/+ fish (**Figure 3B**, upper). After electrophoresis through a 3% agarose gel, two RT-PCR products were observed from null/+ fish: one corresponding to transcripts of the wild type allele (290bp) and the other corresponding to transcripts of the null allele (274bp). The signal from the smaller RT-PCR product, from the null allele transcript of *sorl1*, was visibly less intense than that of the larger, wild type transcript RT-PCR product.

Additionally, the wild type RT-PCR product from null/+ brains appeared less intense than the wild type RT-PCR product from +/+ brains (**Figure 3B**, lower). Although this method of analysis is not strictly quantitative, these results support that the *sorl1* null allele transcript is likely subject to NMD and that there are fewer copies of the wild type transcript in the null/+ brains relative to their +/+ siblings. In summary, the mutant alleles of *sorl1* are likely subject to NMD.

### Transcriptome analyses of null/+, EOfAD-like/+ and transheterozygous mutant brains compared to +/+ sibling brains

Do dominant EOfAD mutations of *SORL1* act through loss- or gain-of-function mechanisms or both? To address this question, we sought to make a detailed molecular comparison of the effects of our EOfAD-like mutation and our putatively null mutation. We performed RNA-seq on brain-derived mRNA in a “four-way” analysis (**Figure 1**) with the aim of identifying the global changes to gene expression caused by mutations in *sorl1*. We first visualised the relationship between the individual samples by principle component analysis (PCA) (**Additional File 4**). We observed limited separation of samples across principal component 2 (PC2), which explained only ~6% of the total variance in this dataset. This suggests that *sorl1* genotype had only subtle effects on the brain transcriptomes. The PCA also revealed that the first principal component, PC1, the largest source of variation in this dataset, appeared to be highly correlated with the library size of the samples. Therefore, to assist with identifying the effects due to *sorl1* genotype, we adjusted for this variation by using the negative control gene (RUVg) method from RUVSeq. An additional PCA on the RUVSeq-adjusted logCPMs (log_2_ counts per million) showed that samples now mostly separated by genotype across PC1, although some variability was observed within the wild type and transheterozygous mutant samples. Importantly, after RUVSeq, PC1 was no longer dependent on library size (**Additional File 4**).

### Mutations in *sorl1* have subtle effects on gene expression in the brain

To our knowledge, an *in vivo* characterisation of the effects of mutations in *sorl1* on the brain transcriptome has not previously been performed. Therefore, we investigated which genes were dysregulated due to heterozygosity for the EOfAD-like mutation, or the null mutation, or complete loss of wild type *sorl1* function (i.e. in the transheterozygous mutant brains).

We could detect differential expression of only one gene, *cytochrome c oxidase subunit 7A1* (*cox7a1*) in EOfAD-like/+ brains relative to wild type brains. In the null/+ brains, we detected 15 differentially expressed (DE) genes relative to wild type, while transheterozygosity for the EOfAD-like and null *sorl1* mutations revealed 20 DE genes relative to wild type (**Figure 4A** and **B**, **Additional File 5)**. Four DE genes were found to be shared between the null/+ and transheterozygous mutant brains: *pim proto-oncogene*, *serine/threonine kinase related 191*, (*pimr191*), *cue domain containing 1b*, (*cuedc1b*), *CABZ01061592.2* and *heat shock 105/110 protein 1* (*hsph1*). Differential expression of *cox7a1* was observed in both the EOfAD-like/+ and transheterozygous mutant samples. No DE genes were shared between the null/+ and EOfAD-like/+ brains relative to their wild type siblings, suggesting that, at the single-gene level, the EOfAD-like mutation does not act through a simple loss-of-function mechanism (**Figure 4C**).

**Figure 4:**
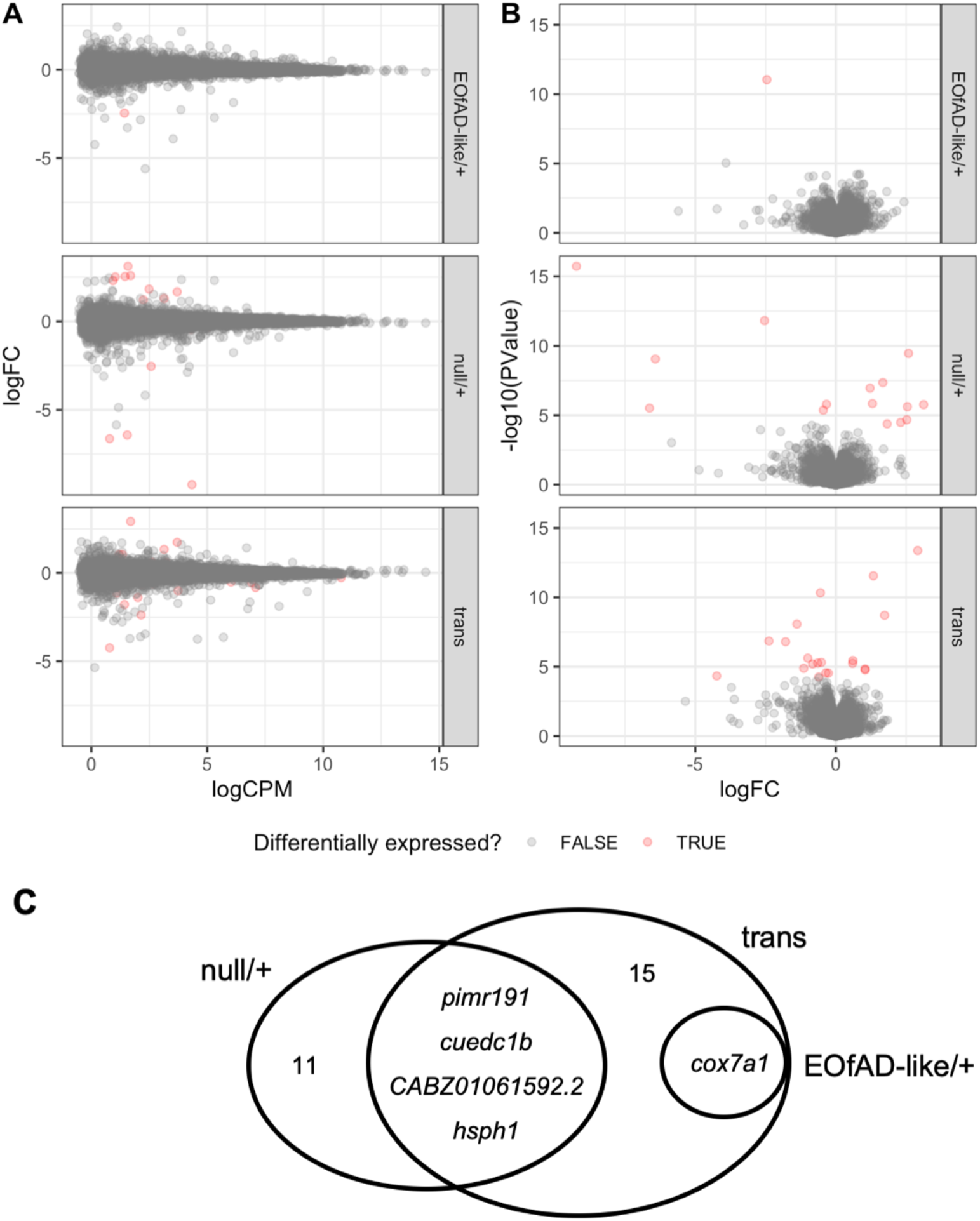
Differential gene expression analysis **A.** Mean-difference (MD) plots for each comparison between the three *sorl1* mutant samples and the wild type samples. **B.** Volcano plots for each comparison between the three *sorl1* mutant samples and the wild type samples. **C**. Venn diagram displaying the overlap of differentially expressed genes in each comparison. “trans” denotes transheterozygous.

### Gene set enrichment analysis reveals that an EOfAD-like mutation in *sorl1* appears to have both loss-of-function and gain-of-function properties

Since few DE genes were observed in the *sorl1* mutant brain transcriptomes relative to the wild type brain transcriptomes, we aimed to obtain a more complete view of the changes to gene expression and cellular function by performing enrichment analysis on the entire list of detectable genes in the RNA-seq experiment. If EOfAD-like mutations in *sorl1* caused, exclusively, loss-of-function effects, we would expect similar gene sets to be altered in the EOfAD-like/+ brains and the null/+ brains.

For enrichment analysis, we used the KEGG and HALLMARK gene sets from MSigDB. The HALLMARK gene sets represent 50 distinct biological processes built both computationally and manually from the intersections between several gene set collections. These gene sets are useful to generate an overall view of key, distinct biological processes (38). In contrast, the KEGG gene sets (186 gene sets) give a more precise view of changes to gene expression in particular cellular processes (39), but pathways are more likely to share many common genes. We used these gene sets in the self-contained gene set testing method fry (43), and the competitive gene set testing method camera (44). However, we did not find any statistical evidence that any of the KEGG or HALLMARK gene sets were significantly altered in any comparison (for the top 10 most altered pathways, see **Additional File 6**).

We also used fgsea (47), which is the fast implementation of the self-contained gene set testing method GSEA (46). Using fgsea, we observed gene sets significantly showing changes as a group after Bonferroni adjustment for multiple testing in each comparison (**Additional File 6**). However, fgsea does not take into account inter-gene correlations and can be prone to false positives (46). Therefore, we a calculated a consensus p-value derived from the p-values from each of the three methods of enrichment analysis by calculating the harmonic mean p-value (48). After FDR adjustment of the harmonic mean p-values, we found 11 gene sets to be significantly altered by heterozygosity for the null mutation, 16 gene sets significantly altered by heterozygosity for the EOfAD-like mutation and 11 gene sets significantly altered by transheterozygosity for the null and EOfAD-like mutations. A summary of the significant gene sets is shown in **Figure 5**.

**Figure 5:**
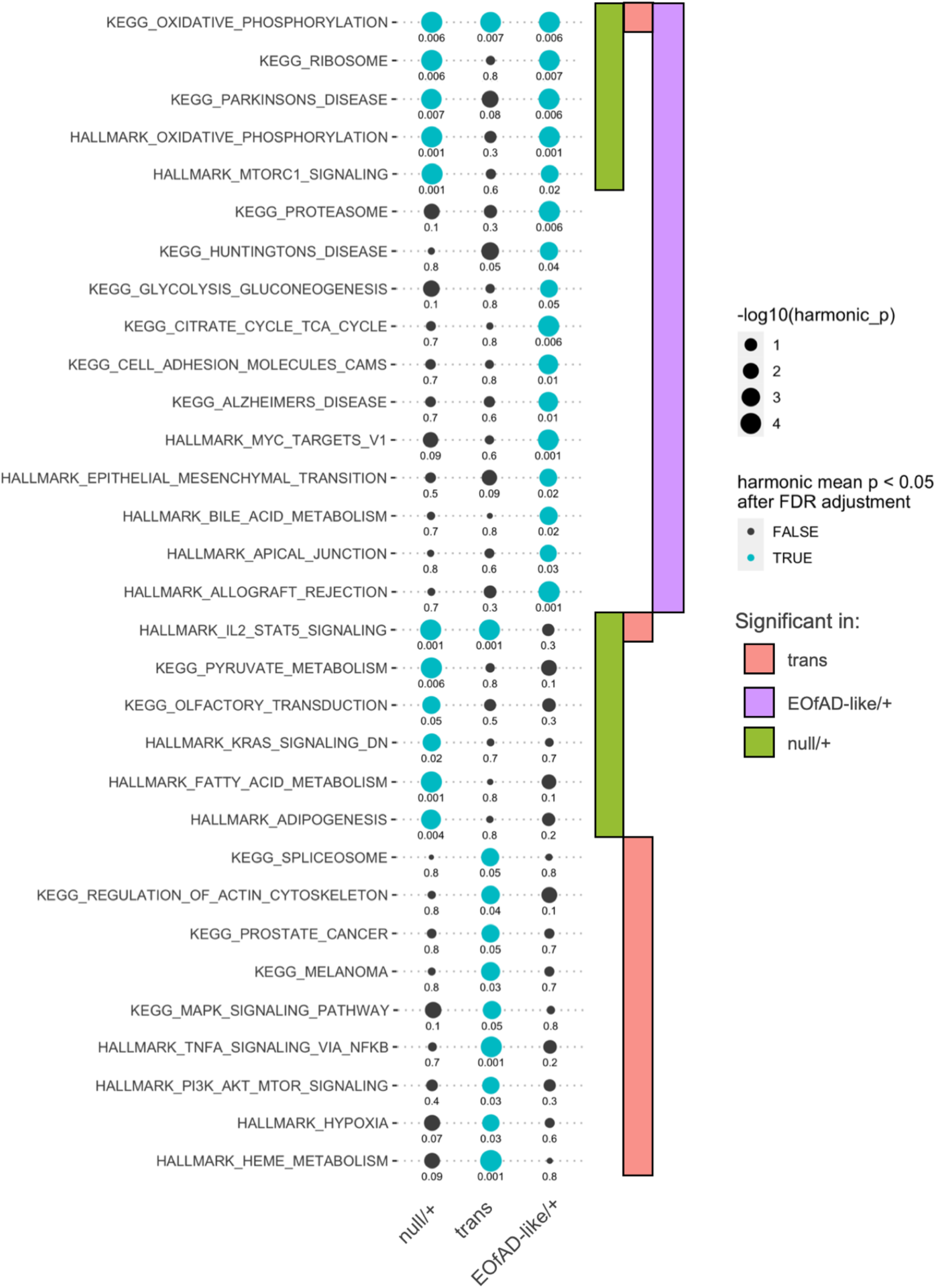
Gene set enrichment analysis Figure 5 depicts the significantly altered KEGG and HALLMARK gene sets in the null/+, EOfAD-like/+ and transheterozygous mutant brains relative their wild type siblings. The sizes of the dots indicate the negative log_10_ of the harmonic mean p-value (i.e. larger dots indicate greater statistical significance) of three methods of enrichment analysis (fry, camera and fgsea) when combined within each comparison. Gene sets are grouped as to the comparisons in which they are significant. Numbers associated with each dot are FDR adjusted harmonic mean p-values.

Out of the 16 gene sets significant in the EOfAD-like/+ brains, 5 were also observed to be significantly altered in the null/+ brains. Intriguingly, three of these gene sets involve mitochondria (the KEGG and HALLMARK gene sets for oxidative phosphorylation, and Parkinson’s disease). Genes encoding ribosomal subunits, and genes involved in the mammalian target of rapamycin complex 1 (mTORC1) signalling pathway, also appear to be affected by the EOfAD-like mutation and the null mutation, indicating that these effects are likely due to decreased *sorl1* function.

Intriguingly, heterozygosity for the EOfAD-like mutation also gives rise to changes in gene expression which do not appear to be affected in the null/+ brains, suggesting gain-of-function action. Interestingly, the KEGG gene set for Alzheimer’s disease is one of these significantly altered gene sets. Other gene sets span inflammation, cell adhesion, protein degradation, bile acid metabolism, myc signalling and the tricarboxylic acid (TCA) cycle.

Many HALLMARK and KEGG gene sets include shared genes and so are commonly co-identified as significantly altered in enrichment analysis. Therefore, we inspected whether any of the gene sets found to be significantly altered were being driven by changes to expression of the same genes. Genes in the “leading edge” of the fgsea algorithm can be interpreted as the core genes which drive the enrichment of the gene set. We found the leading edge genes within the significant gene sets in each *sorl1* genotype comparison to be mostly independent of one another (**Additional File 7**). However, we observed some overlap of leading edge genes for the oxidative phosphorylation and neurogenerative diseases gene sets in the EOfAD-like/+ and null/+ comparisons (i.e. KEGG_ALZHEIMERS_DISEASE, KEGG_HUNTINGTONS_DISEASE and KEGG_PARKINSONS_DISEASE). This indicates that the enrichment of these gene sets is driven mostly by the same gene differential expression signal (genes encoding electron transport chain components) (**Additional File 7**).

### Changes in gene expression due to *sorl1* mutations are not due to broad changes in cell-type distribution

One possible, artifactual explanation for the apparent changes in gene expression we observe could be changes to the proportions of cell-types within the brains of fish with different *sorl1* genotypes. To account for this, we obtained a list of representative marker genes for four broad cell types found in zebrafish brains: neuron, astrocyte and oligodendrocyte markers (52), and microglial markers (53). We then examined their expression across the brain samples. We did not observe any obvious differences between genotypes, supporting that the changes seen in gene expression are not due to broad changes in cell type proportions within the brains (**Additional File 8**).

### The effects of heterozygosity for an EOfAD-like mutation are distinct from those of complete loss of wild type *sorl1* function

In research in AD genetics, it is common for cell- and animal-based studies to analyse mutations in their homozygous state. This allows easier identification at contrasting functions between mutant and wild type alleles. However, this ignores that, in humans, EOfAD mutations and LOAD risk variants are rarely homozygous and that interaction between mutant and wild type alleles can generate unique molecular effects. The transheterozygous genotype in our study can be considered somewhat similar to a homozygous loss-of-function, since it lacks the wild type *sorl1* allele. We found that genes of the oxidative phosphorylation pathway were the only genes to be significantly affected as a group in common across the EOfAD-like/+, null/+, and transheterozygous mutant brains (**Figure 5**). Concerningly, the genes which make up this gene set mostly showed opposite direction of change in the EOfAD-like/+ (and the null/+ brains) compared to the transheterozygous mutant brains (**Figure 6A, Additional File 9**). Indeed, we found that the majority of genes within the gene sets significantly altered in the EOfAD-like/+ brains generally showed opposite directions of change in the transheterozygous mutant brains (**Figure 6**). This suggests that heterozygosity for this EOfAD-like mutation has very different effects to complete loss of normal *sorl1* function, and highlights the importance of analysing genetic models that mimic the heterozygous genetic state of the human disease.

**Figure 6:**
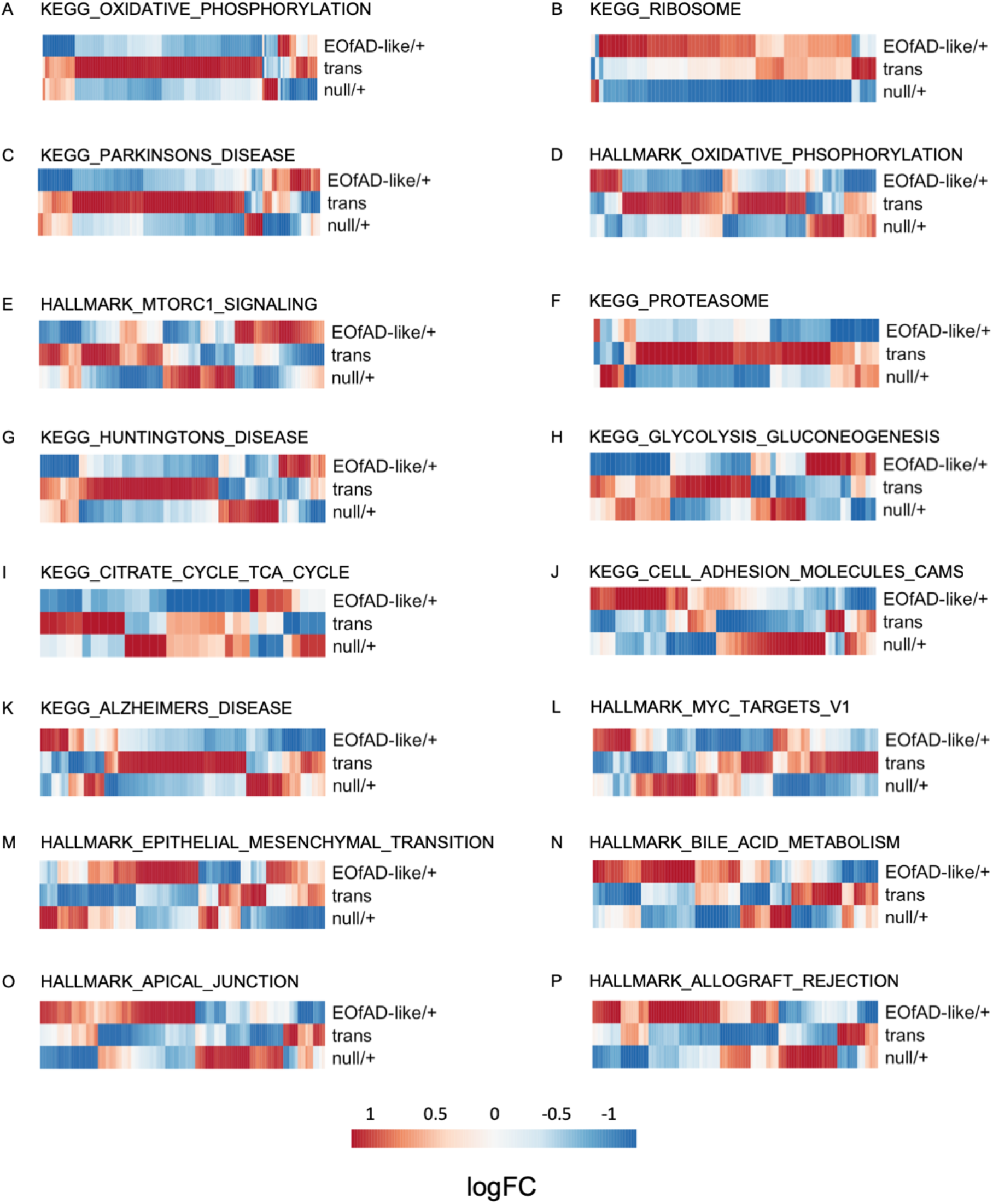
Heterozygosity for an EOfAD-like mutation in sorl1 often results in changes to gene expression in the opposite direction to more complete loss of wild type sorl1 function. **A-P.** LogFC of the genes making up the gene sets in each comparison of *sorl1* mutants. Only the gene sets found to be significantly enriched in EOfAD-like/+ brains are shown. The colour legend indicates the magnitude of the logFC in the heatmaps (red is upregulation, blue is downregulation).

### Loss of *sorl1* function may cause iron dyshomeostasis

A recent, sophisticated study in (54) demonstrated that impairment of acidification of the endolysosomal system inhibits the ability of cells to reduce ferric iron to more reactive ferrous iron. This results in a pseudo-hypoxic response, since a master regulator protein of the cellular response to hypoxia, HIF1-α, is degraded under normal oxygen conditions by a ferrous iron-dependent mechanism. Ferrous iron deficiency was shown to result in mitochondrial dysfunction and inflammation. We recently developed a method to detect possible iron dyshomeostasis in RNA-seq data, by examining changes to the expression of genes with iron-responsive elements (IREs) in untranslated regions (UTRs) of their mRNAs (41). We hypothesised that the changes in gene expression observed in the *sorl1* mutant brains in the oxidative phosphorylation pathway may be co-occurring with changes in, or responses to, cellular iron levels. To test this, we obtained the IRE gene sets developed in (41). These four gene sets consist of genes classified as having a canonical or non-canonical IRE in the UTRs of their mRNAs, and having an IRE motif in the 5’ or 3’ UTR of the mRNA. We performed an enrichment analysis similar to those performed above for the KEGG and HALLMARK gene sets to determine whether each set of IRE genes showed changes in expression as a group. We found that gene transcripts containing 3’ IREs were significantly altered in the transheterozygous mutant brains. Expression of gene transcripts with both canonical and non-canonical 3’ IREs generally was decreased. We did not find any statistical evidence that any of the other *sorl1* genotypes affected the expression of gene transcripts containing IREs (**Figure 7**).

**Figure 7:**
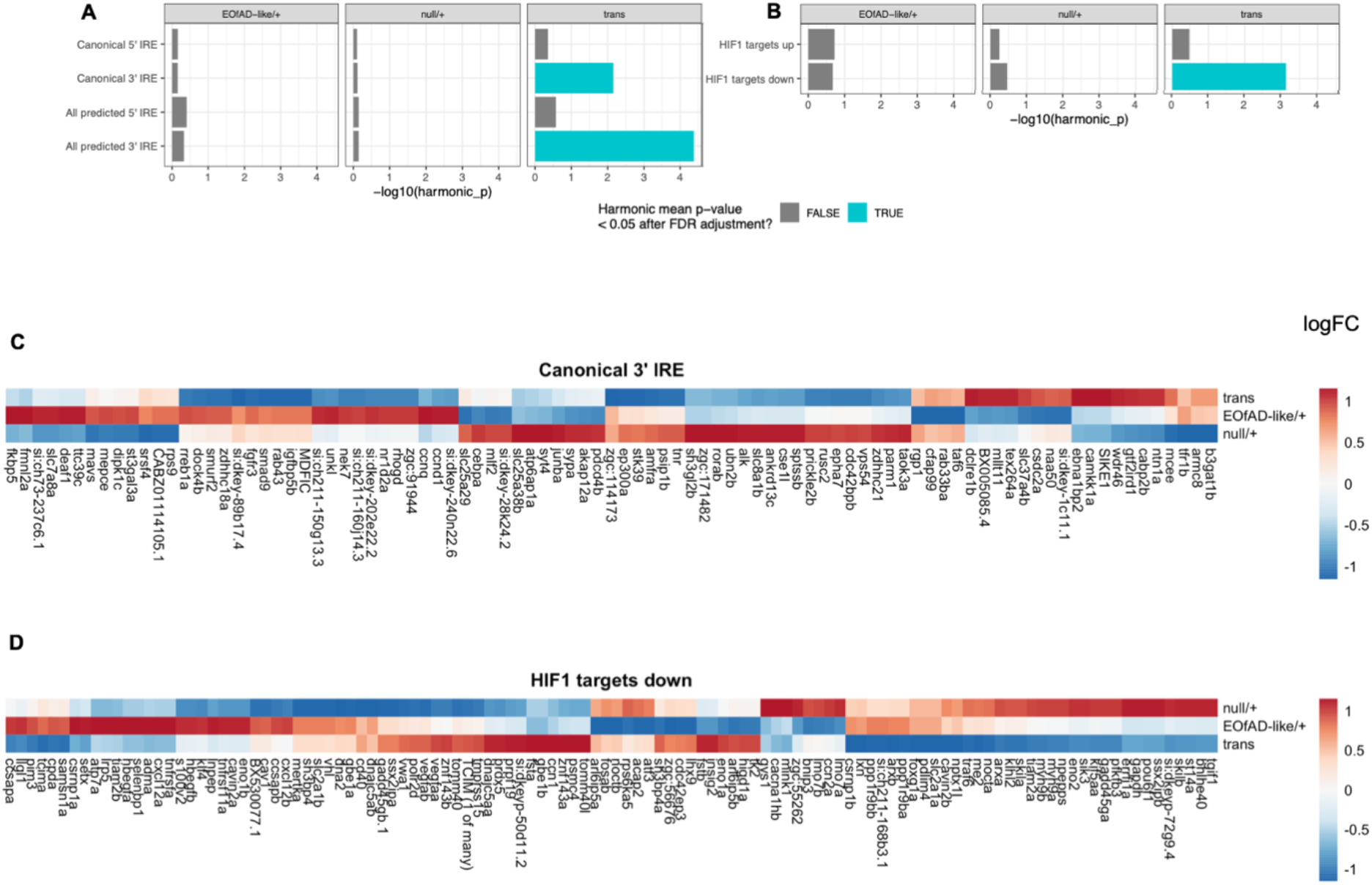
Loss of wild type *sorl1* results in dysregulation of genes with an iron response element in their transcript’s 3’ untranslated region, and of genes involved in the cellular response to hypoxia. The harmonic mean p-values for the enrichment analyses of gene sets with **A**, iron response elements (IRE) in their untranslated regions, and **B**, regulated by the HIF1 transcription factor, are shown with the significant gene sets in blue. The logFCs of genes in brains of each genotype with **C**, a canonical IRE in their transcript’s 3’ untranslated region and **D**, shown to be downregulated by loss of HIF1 activity. The genes and comparisons are clustered based on their Euclidian distance.

As mentioned earlier, changes to iron homeostasis result in stabilisation of protein HIF1-α. This combines with the protein HIF1-β to form the transcription factor HIF1 that then initiates a pseudo-hypoxic response (54). The HALLMARK gene set for hypoxia was found to be significantly altered in the in transheterozygous mutant brains, but not in the other *sorl1* genotypes. We aimed to characterise further whether the *sorl1* mutant fish were responding transcriptionally in a hypoxia-like manner by performing an enrichment analysis on the HIF1 target gene sets GROSS_HYPOXIA_VIA_HIF1A_UP and GROSS_HYPOXIA_VIA_HIF1A_DN (hereafter referred to as HIF1 targets). These gene sets were described in (42) and show the genes found to be up- and down-regulated in response to siRNA knockdown of HIF1-α respectively. We found that the downregulated HIF1 targets were significantly altered in the transheterozygous samples, but not in any other mutant samples. However, downregulation of the genes of the GROSS_HYPOXIA_VIA_HIF1A_DN gene set in transheterozygous mutant brains was not consistently observed (**Figure 7D**). Together, these results support that complete loss of wild type *sorl1* results in changes to iron homeostasis.

## Discussion

Here, we generated and analysed the first knock-in (endogenous gene) animal models of null and EOfAD-like mutations in *SORL1.* We introduced an EOfAD-like mutation into the zebrafish orthologue of *SORL1* using TALENs. This mutation, V1482Afs, models the human mutation C1478* in *SORL1.* C1478* was identified in a French family and appears to segregate with AD with a dominant inheritance pattern (16). The C1478* transcript of *SORL1* was shown to be likely subject to NMD in peripheral blood (16). We showed that the zebrafish V1482fs transcript of *sorl1* also appears to be subject to NMD, as there is a clear imbalance in the expression of the mutant and wild type alleles of *sorl1* present in the EOfAD-like/+ brains. Nevertheless, transcripts of the EOfAD-like allele can still be detected, and so, potentially, can be translated to produce a truncated protein that may act in a dominant manner.

We aimed to identify the global changes to gene expression in a relatively unbiased manner in the brains of young adult (6 month old) zebrafish, heterozygous for the EOfAD-like mutation. At 6 months of age, zebrafish are recently sexually mature, and we consider this to be the equivalent of early adulthood in humans. Analysis at this age allowed us to model the early changes to gene expression occurring in EOfAD mutation carrier brains long before cognitive symptoms are predicted to occur. We also wanted to investigate whether the EOfAD-like mutation was deleterious due to haploinsufficiency or neomorphism. Therefore, for comparison, we generated a putatively null mutation in *sorl1,* R122Pfs, which is predicted to encode a protein product lacking any of the functional domains of wild type Sorl1 protein. We generated a family of zebrafish composed of sibling fish heterozygous for the EOfAD-like mutation, or heterozygous for the null mutation, as well as fish transheterozygous for the null and EOfAD-like mutations and their wild type siblings (Figure 1). We initially intended to omit transheterozygous fish from our analysis, as this genotype is unrepresentative of the human EOfAD genetic state. However, recently, a human patient transheterozygous for *sorl1* mutations was described in the literature (55). Also, the transheterozygous fish lack wild type *sorl1* and, therefore, might be informative as an extreme loss-of-function phenotype. A family of siblings was raised together in a similar environment (in three tanks side-by-side in a re-circulating water system) to reduce genetic and environmental sources of variation in our analysis thereby maximising the possibility of identifying subtle changes in gene expression due to *sorl1* genotype.

### Mutations in *sorl1* have subtle effects on the young-adult brain transcriptome

We identified a small number of DE genes in the brains of each of the different *sorl1* mutant fish relative to their wild type siblings (**Figure 4**). The transheterozygous mutant fish had the greatest number of genes identified as DE. This was not surprising as these fish lack any wild type *sorl1* expression. Only *cox7a1* was found to be significantly differentially expressed in the EOfAD-like/+ brains. This nuclear gene encodes a subunit of the mitochondrial cytochrome c oxidase complex, the terminal component of the respiratory electron transport chain. In humans, this gene has two isoforms, one which is mainly expressed in the heart and skeletal muscle and the other which is mostly expressed in the brain and uterus (56). Only one of the three isoforms of *cox7a1* known to exist in zebrafish was expressed in the brains of these fish (data not shown).

In comparison to heterozygosity for the EOfAD-like mutation in *sorl1*, heterozygosity for an EOfAD-like mutation in *psen1*, the gene most commonly mutated in EOfAD (14, 57), resulted in differential expression of 251 genes in young-adult, zebrafish brains (58). This suggests that EOfAD-like mutations in *sorl1* are not as disruptive to cell function as EOfAD-like mutations in *psen1.* Campion, Charbonnier (23) noted that carriers of the C1478* mutation generally had an age of onset closer to the established LOAD threshold of 65 years, which arbitrarily defines the difference between EOAD and LOAD, and that unaffected, aged (≥ 66 years of age), carriers of this mutation exist. This, together with our results, supports the idea that *SORL1* resembles more closely a LOAD genetic risk locus, rather than an EOfAD-causative locus such as *PSEN1.*

### EOfAD-like mutations in *sorl1* appear to have both haploinsufficient loss-of-function and neomorphic gain-of-function properties

EOfAD mutations in the *PSENs* and *APP* follow a “reading frame preservation rule”, where mutations preventing translation of C-terminal sequences of the protein do not cause AD (reviewed in (15)). Reading frame-truncating mutations in *SORL1* have been found in EOfAD families so EOfAD mutations in *SORL1* do not follow this rule (16, 17). Since the protein-truncating mutations have been shown to be subject to NMD (16), it is thought that these mutations have loss-of-function disease phenotypes due to haploinsufficiency.

Understanding the mechanism behind the action of EOfAD mutations in *SORL1* is critical to understanding how these mutations increase risk for developing the disease. We observed that heterozygosity for the EOfAD-like mutation resulted in changes to gene expression involved in cellular processes including energy production, mRNA translation and mTORC1 signalling. These changes are also observed in null/+ brains, supporting that they are loss-of-function effects due to haploinsuffiency. Restoring *sorl1* expression levels in EOfAD mutation carrier brains could provide a means of ameliorating these apparently pathological changes. We also observed significant changes to gene expression involved in cellular processes in EOfAD-like/+ brains which are not significantly altered in the null/+ brains, suggesting that these effects are due to neomorphic gain-of-function. These cellular processes are likely affected by the residual expression (after NMD) of N-terminal Sorl1 protein domains without the C-terminal sequences. These effects span other pathways involved in energy production, such as glycolysis and the TCA cycle, cell adhesion, myc signalling, protein degradation and cell adhesion. However, we cannot rule out that these changes are not occurring in the null/+ brains, as the analysis of the null/+ brains had less statistical power than for the other genotypes (n = 4 rather than n = 6). For example, the myc targets and glycolysis gene sets had FDR adjusted p-values of approximately 0.1 in the null/+ brains. With n = 6, these gene sets may have been found to be statistically significantly altered.

### Cellular processes potentially dependent on normal *sorl1* function

Our enrichment analysis has identified cellular processes not previously known to require normal *sorl1* function. We observed gene sets linked to energy production to be significantly altered in *sorl1* mutant brains relative to their wild type siblings. Genes involved in the oxidative phosphorylation pathway were observed to have altered expression as a group in all three *sorl1* mutant genotypes (**Figure 5**). Expression of genes in this pathway were mostly reduced in the EOfAD-like/+ and the null/+ brains, suggesting that this effect is due to loss-of-function through haploinsufficiency. Intriguingly, expression of genes in the oxidative phosphorylation pathway was mostly increased in the transheterozygous mutant brains relative to their wild type siblings (**Additional File 9**). This opposite direction of change was also observed in other gene sets involving energy production (i.e. glycolysis in **Figure 6H** and the TCA cycle in **Figure 6I**). These effects are subtle, as the genes individually were not detected as differentially expressed (other than *cox7a1*) and the signal was only detected at the pathway level. Genes encoding the components of the electron transport chain have been shown previously to be dysregulated in early in AD (59–61), supporting that changes to mitochondrial function are an early pathological change in AD.

One explanation for changes in gene expression in the oxidative phosphorylation pathway could be changes in the function of the mitochondrial associated membranes (MAMs) of the endoplasmic reticulum. MAMs are lipid-raft like regions of the endoplasmic reticulum which form close associations with mitochondria and that regulate mitochondrial activity through calcium ion release (reviewed in (62–64)). It has been shown that the presenilins, APP, BACE1 (β-secretase) and other components of the *γ*-secretase complex are enriched in MAMs relative to the rest of the endoplasmic reticulum (65, 66). SORL1 has been shown to form a complex with APP and BACE1 (67), and is also cleaved by *γ*-secretase (68). We have shown previously that SORL1 is present in the MAM in the brains of mice (24) (see **Additional File 10** for a reproduction of this result). SORL1 may bind APP in the MAM and affect its cleavage by BACE1 and *γ*-secretase (which can function as a supramolecular complex (69)). Whether mutations in *sorl1* affect normal MAM function is yet to be determined.

We also found evidence that iron homeostasis is affected by complete loss of wild type *sorl1* function. The transcripts of genes possessing a canonical or non-canonical IRE in their 3’ UTR were found to be dysregulated in transheterozygous mutant brains. It has generally been assumed that transcripts with an IRE in their 3’ UTR have increased stability under cellular iron deficiency, and decreased stability under iron overload (reviewed in (12)). However, this is not necessarily the case and transcripts with 3’ IREs can have increased or decreased stability under cellular iron deficiency (41). Therefore, whether transheterozygous mutant brains are under cellular iron deficiency or overload is still unclear and further investigation is warranted.

The cellular response to hypoxia is mediated by hypoxia-inducible factor 1, α subunit (HIF1-α) in an iron dependent manner (HIF1-α is less stable under normoxia and in ferrous iron replete conditions). We also observed that genes involved in the cellular response to hypoxia are affected in transheterozygous mutant brains. Since oxygen and iron homeostasis are both involved in the stabilisation of HIF1-α, this further supports a role for *sorl1* in iron homeostasis. Yambire, Rostosky (54), showed that insufficient acidification of the endolysosomal system results in a pseudo-hypoxic response (stabilisation of the HIF1-α not due to a low oxygen environment) and the consequent ferrous iron deficiency led to mitochondrial instability and an increase in markers of inflammation. We also observed that genes involved in inflammation (HALLMARK_TNFA_SIGNALING_VIA_NFKB and HALLMARK_IL2_STAT5_SIGNALING gene sets) are significantly affected as a group in the transheterozygous mutant brains consistent with a role of *sorl1* function in iron homeostasis. Since the effects on iron homeostasis, hypoxic response and inflammation are most evident when both copies of wild type *sorl1* are lost, this implies that these effects may be recessive in nature.

Recently, Jiang and colleagues (70), demonstrated that increased expression of the β-CTF/C99 fragment of APP (produced by β-secretase cleavage, also known as “amyloidogenic processing” of APP) results in decreased acidification of the endolysosomal system in cells. SORL1 physically interacts with APP to modulate its amyloidogenic processing (71–73), and this suggests a molecular mechanism for SORL1’s action in iron homeostasis. A more indirect action on iron homeostasis via modulation of APP’s C99 fragment would be consistent with the generally later age of AD onset seen for carries of putative EOfAD mutations in *SORL1* compared to carriers of EOfAD mutations in *APP* (16).

Surprisingly, we did not observe any endolysosomal gene sets to be significantly altered by mutation of *sorl1* mutant genotype despite that one of the primary sites of action of SORL1 protein is thought to be within the endolysosomal system (71). Loss of *SORL1* by CRISPR-Cas9 mutagenesis in neurons (but not in microglia) derived from human induced pluripotent stem cells (hiPSCs) resulted in early endosome enlargement (74), a commonly observed pathology in AD brains (75, 76). To determine whether endolysosomal defects occur in our *sorl1* mutant zebrafish brains requires further investigation, as any effects may be too subtle to detect when analysing RNA-seq data from whole brains, or the effects may not be present until later ages.

### Heterozygosity for an EOfAD-like mutation appears to have effects distinct from complete loss of *sorl1* function

Our results support that caution should be used when assuming that homozygous disease mutation model animals have similar, only more severe, phenotypes compared to heterozygous animals. We found little similarity between the transcriptomes of EOfAD-like/+ and transheterozygous mutant brains. At the pathway level, most processes affected in the transheterozygous mutant brains appeared unaffected in the EOfAD-like/+ mutant brains and vice versa. Only the oxidative phosphorylation pathway was affected in both comparisons. However, as discussed previously, the overall changes in oxidative phosphorylation gene expression appeared to be in opposite directions in the two genotypes, suggesting different mechanisms of action. This highlights the importance of using a genetic model closely resembling the genetic state of the human disease to make exploratory analyses of the effects of a human mutation in an unbiased manner. Hargis and Blalock (13), showed that some of the commonly used transgenic “mouse models of AD” (which do not closely reflect the genetic state of AD) have very little transcriptome concordance with human LOAD, or even with each other. We recently demonstrated little similarity between brain transcriptome changes in young adults of the popular 5xFAD transgenic mouse model and zebrafish heterozygous for an EOfAD-like, knock-in mutation of *psen1* (41). The similar apparent effects of EOfAD-like mutations in *psen1* and in *sorl1* on oxidative phosphorylation in the young adult brains of zebrafish supports that knock-in models of EOfAD mutations may provide more consistent modelling of the early molecular pathogenesis of EOfAD.

Transcriptomic analyses of knock-in models of EOfAD mutations are beginning to be appear in the literature. Our analysis of an EOfAD-like mutation in the zebrafish orthologue of *PSEN1* found that heterozygosity for this mutation affects expression of genes involved in acidification of the endolysosomal system, oxidative phosphorylation and iron homeostasis in the brain (41, 58). Additionally, male knock-in *APOE* mice, which carry a humanised form of *APOE* (with genotype *APOE ε3/ε4*) associated with LOAD, showed changes to brain gene expression in the oxidative phosphorylation pathway at 15 months of age (77). Interestingly, these mice (and *APOE ε4/ε4* mice) also show abnormalities in the endolysosomal system, which could mean that iron homeostasis is dysregulated. These results together suggest that iron dyshomeostasis and changes in oxidative phosphorylation may be unifying characteristics of AD, linking the rare, early-onset, familial subtype of AD with the common, late-onset, sporadic subtype.

### Candidate genes for regulation by the SORL1 intracellular domain in the brain

Finally, it is worth noting that the genes detected as differentially expressed in transheterozygous mutant brains likely include candidates for regulation by the intracellular domain of Sorl1 protein (Sorl1-ICD). It has been shown that human SORL1 protein is cleaved by both α- and *γ*-secretases to produce various fragments (68, 78, 79). The human SORL1-ICD has been demonstrated to enter the nucleus and regulate transcription (78). However, the genes it regulates are yet to be identified. Both the EOfAD-like and null mutations of zebrafish *sorl1* are predicted to truncate Sorl1 protein so that it lacks the Sorl1-ICD sequences. Therefore, the genes identified as differentially expressed in the transheterozygous mutant brains are possible direct targets for regulation by the Sorl1-ICD and warrant further investigation.

## Conclusion

We have made an initial, *in vivo* characterisation of changes in gene expression in the brain due to mutations in *sorl1.* Our results provide insight into novel cellular processes involving *sorl1*, such as energy production and iron homeostasis. Changes in these processes may represent early cellular stresses ultimately driving the development of AD. Our results support both loss- and gain-of-function actions for EOfAD-like mutation of *sorl1*. The severity of *sorl1* mutation effects are more consistent with human SORL1’s action as a LOAD risk locus, since the effects are subtle compared to an EOfAD-like mutation in *psen1.* Future work may include “four-way” analyses of brain transcriptomes at older ages, where any effects may be more pronounced as well as further characterisation of those cellular processes we have identified as altered by mutations in *sorl1*.

## Abbreviations

Aβ: amyloid beta
AD: Alzheimer’s disease
ANOVA: analysis of variance
APOE: apolipoprotein E
APP: amyloid beta a4 precursor protein
BDNF: brain-derived neurotrophic factor
bp: base-pair
Cas9: CRISPR associated protein 9
cDNA: complementary DNA
cox7a1: cytochrome c oxidase subunit 7A1
cpf1: CRISPR associated protein 12a
cpm: counts per million
CRISPR: clustered regularly interspaced short palindromic repeats
cuedc1b: cue domain containing 1b
DE: differentially expressed
DNaseI: deoxyribonuclease I
dqPCR: digital quantitative polymerase chain reaction
EGF: epidermal growth factor
EOfAD: early-onset familial Alzheimer’s disease
FC: fold change
FN: fibronectin
GEO: Gene Expression Omnibus
GSEA: gene set enrichment analysis
HIF1-α: hypoxia-inducible factor 1-alpha
hiPSCs: human induced pluripotent stem cells
ICD: intracellular domain
IRE: iron-responsive element
KEGG: Kyoto Encyclopedia of Genes and Genomes
LDLR: low density lipoprotein receptor
LOAD: late-onset Alzheimer’s disease
MAM: mitochondrial-associated membrane
mRNA: messenger RNA
MSigDB: molecular signatures database
mTORC: mammalian target of rapamycin complex
NMD: nonsense mediated mRNA-decay
nt: nucleotide
PAM: protospacer adjacent motif
PC: principal component
PCA: principal component analysis
PCR: polymerase chain reaction
PSEN: presenilin
RNA-seq: RNA sequencing
RT-PCRs: reverse transcription polymerase chain reactions
SAHMRI: South Australian Health and Medical Research Institute
siRNA: small interfering RNA
SORL1: sortilin-related receptor 1
TALEN: transcription activator-like effector nucleases
TCA: tricarboxylic acid
TMD: transmembrane domain
trans: transheterozygous
UTR: untranslated region
VPS10: vacuolar protein sorting 10
WT: wild type.

## Declarations

### Ethics approval and consent to participate

All zebrafish work was conducted under the auspices of the University of Adelaide Animal Ethics Committee (permit numbers: S-2017-073 and S-2017-089) and Institutional Biosafety Committee (permit number 15037).

### Consent for publication

Not applicable.

### Availability of data and materials

The raw fastq files and the output from catchKallisto (i.e. the raw transcript abundances) have been deposited in Gene Expression Omnibus database (GEO) under Accession Number GSE151999 available at https://www.ncbi.nlm.nih.gov/geo/query/acc.cgi?acc=GSE151999. All code to reproduce the RNA-seq analysis can be found at https://github.com/karissa-b/sorl1_4way_RNASeq_6m.

### Authors’ contributions

KB performed all experimental and bioinformatic analysis and drafted the manuscript. SMP supervised and provided advice on the bioinformatic analysis and MN and ML supervised all zebrafish work. All authors read and contributed to the final manuscript.

## Acknowledgements

The authors would like to acknowledge Seyyed Hani Moussavi Nik for obtaining part of the funding for this work and Nhi Hin for providing the sets of zebrafish genes containing iron-responsive elements.

## List of Additional Files

**Additional File 1: Genome editing of zebrafish *sorl1*.**

*Additional File 1.pdf*

**A** and **B** show sections of the *sorl1* genomic sequence (ENSG00000137642) with the sense sequence (upper), translation (middle) and anti-sense sequence (lower). **A**. *sorl1* exon 2, with the crRNA binding site and PAM sequence for cleavage by Cpf1 indicated by pink arrows, and the R122 site indicated by the orange bar. **B**. *sorl1* exon 32, with the C1481 codon indicated by the orange arrow and the TALEN binding sites by blue arrows.

**Additional File 2: Generation of mutant zebrafish lines**

*Additional File 2.docx*

A detailed description of the isolation of the mutant lines of zebrafish.

**Additional File 3: No observed bias for differential expression with GC content or length.**

*Additional File 3.pdf*

Plots showing a ranking metric using the sign of logFC multiplied by −log10 of the p-value against A) the GC content of the gene and B) the length of the gene. The blue curve indicates the line of best fit from a generalised additive model (gam), whilst the black dashed line represents the y = 0 line. Given the gam fit is a nearly horizontal line mostly overlapping y = 0, a significant bias for GC content or length is likely not present in this dataset. The ranking statistic limits were constrained to −10 and 10 for visualisation purposes, and in B, gene length was plotted on the log10 scale.

**Additional File 4: Principal component analysis**

*Additional File 4.pdf*

**A**. Plot of principal component 1 (PC1) against PC2 from a principle component analysis (PCA) of the logCPMs from each sample. **B**. PC1 against the library size per sample. The linear relationship observed between PC1 and library size suggests that the largest source of variability in this dataset is due to library size. **C.** PC1 against PC2 after removal of 1 factor of unwanted variation using RUVSeq (37). **D.** PC1 no longer depends on library size after RUVSeq transformation.

**Additional File 5: Results of differential expression analysis**

*Additional File 5.xlsx*

Each sheet within the spreadsheet gives the entire results of the differential gene expression due to each of the three *sorl1* genotype comparisons with wild type.

**Additional File 6: Results of enrichment testing**

*Additional File 6.docx*

Results of the individual enrichment testing algorithms (fry, camera and fgsea).

**Additional File 7: Results from enrichment analysis are mostly not driven by expression of the same genes.**

*Additional File 7.pdf*

The upset plots show the overlap of the “leading edge” genes from the fgsea algorithm in each of the significant gene sets in **A)** EOfAD-like/+, **B)** null/+, and **C)** transheterozygous mutant brains. Intersections are shown when the leading edge of the gene sets share three or more genes. Overall, the leading edge genes of the significantly altered gene sets are relatively independent of one another. However, genes in the oxidative phosphorylation gene sets and the gene sets for neurodegenerative diseases (Alzheimer’s, Parkinson’s and Huntington’s diseases) all contain genes encoding components of the electron transport chain and are capturing a portion of the same gene expression signal, which is shown in **D)** and **E)**. Missing positions in **D)** and **E)** indicate that the gene was not in the leading edge for that gene set.

**Additional File 8: Changes to gene expression in young-adult zebrafish brain transcriptomes are likely not due to altered cell type proportions.**

*Additional File 8.pdf*

We obtained representative expression markers of neurons (72 genes), oligodendrocytes (100 genes) and astrocytes (44 genes) from (52) and representative expression markers of microglia (533 genes) from (53). The logCPM distributions for these marker genes in each of the samples are similar, supporting that, broadly, the distribution of these cell types is consistent between samples. Data are shown as violin plots, displaying the kernel probability density of the logCPMs, overlaid with boxplots, showing summary statistics, and coloured by genotype.

**Additional File 9: Pathview visualisation of the KEGG oxidative phosphorylation gene set.**

*Additional File 9.pdf*

The logFC of the genes in the KEGG oxidative phosphorylation gene set are shown in each *sorl1* genotype comparison with wild type. Intensity of the colours indicates the magnitude of the logFC, and white indicates that a gene was not detected as expressed in the RNA-seq experiment. Plot was adapted from pathview (80).

**Additional File 10: SORL1 is localized in the MAM in mouse brains.**

*Additional File 10.pdf*

*Note: This figure is reproduced from the Ph.D. thesis of Anne Lim for ease of access.*

Western blot analysis of subcellular fractions of mouse brain cellular membranes. Each subcellular fraction probed with antibodies against **a)** BACE1, **b)** SORL1 and **c)** LRP/LR. **d)** Shows the identity of fractions western immunoblotting using antibodies recognising proteins MTCO1 in mitochondria (Mito), Na+/K+-ATPase in the plasma membrane (PM), IP3R3 in mitochondrial-associated membranes (MAM), and KDEL, predominantly in non-MAM endoplasmic reticulum (ER). Some cross contamination was observed as MTOC1 was detected in both the mitochondria and plasma membrane, and Na+/K+-ATPase in both the mitochondria and plasma membrane.

Lim, A. H. L. (2015). Analysis of the subcellular localization of proteins implicated in Alzheimer’s Disease. <u>Genetics and Evolution</u>, University of Adelaide. **Doctor of Philosophy (PhD):** 235.

