## Supplementary figures and images for "Brain transcriptome analysis reveals subtle effects on mitochondrial function and iron homeostasis of mutations in the *SORL1* gene implicated in early onset familial Alzheimer’s disease"

### Additional File 1

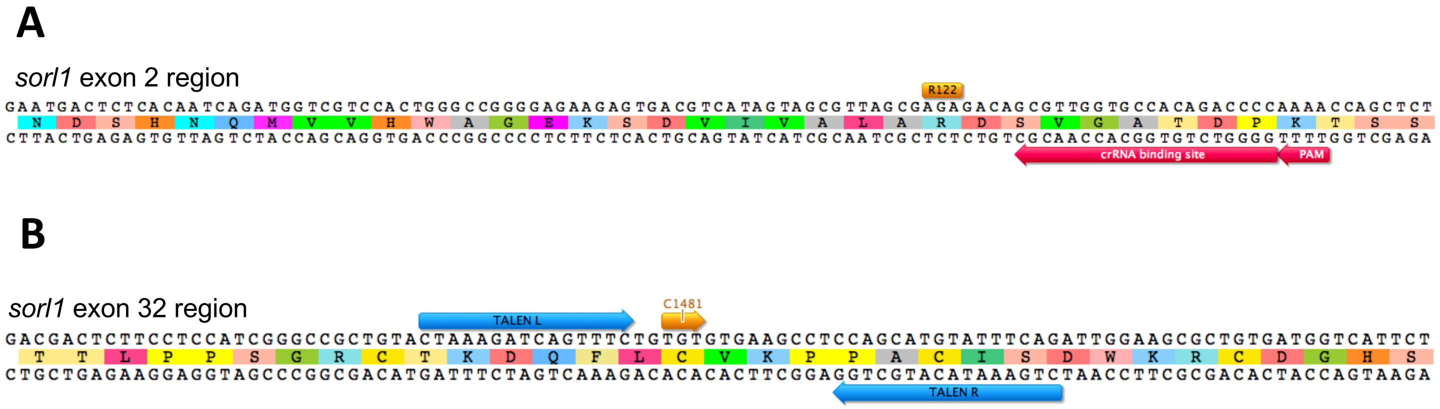

### Additional File 3

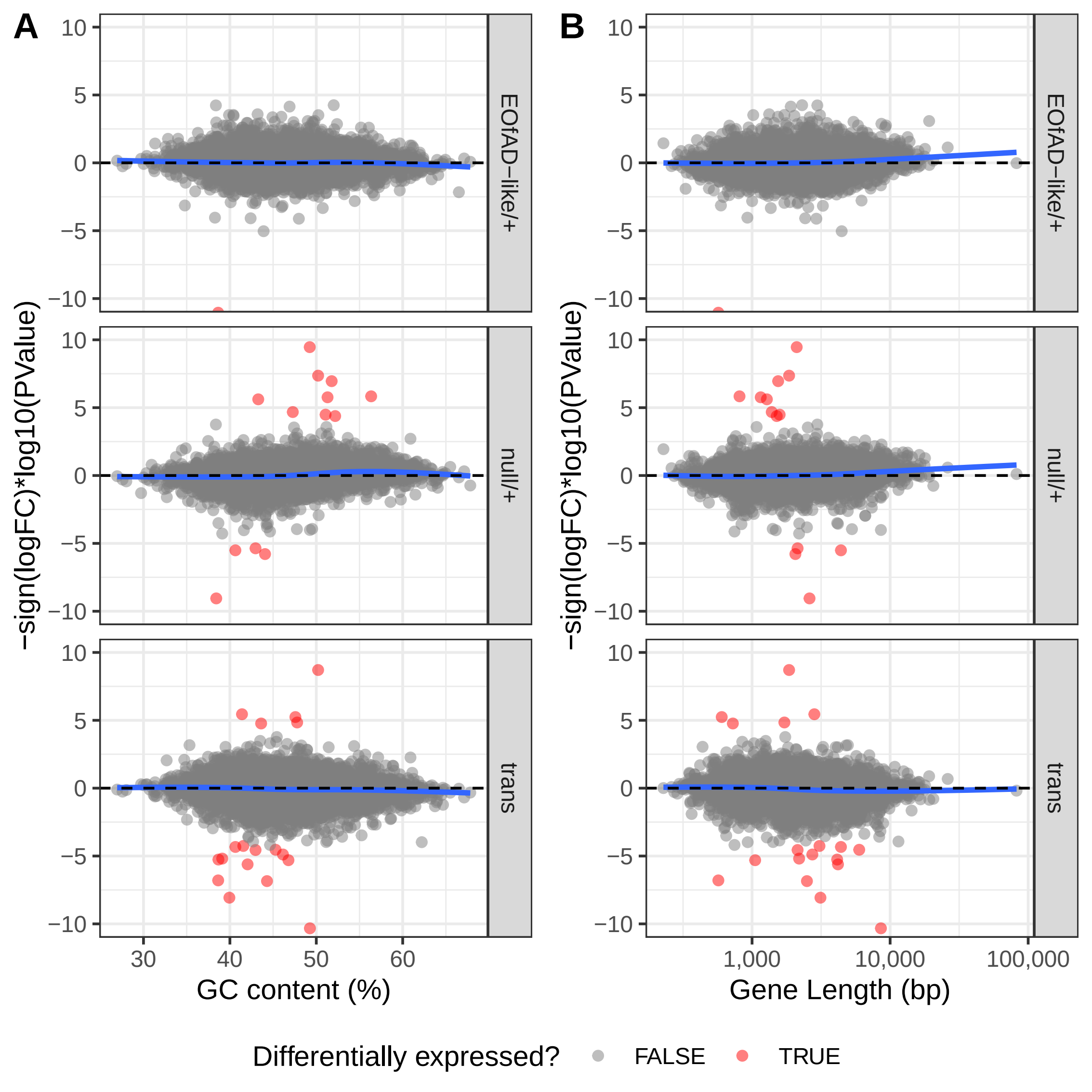

### Additional File 4

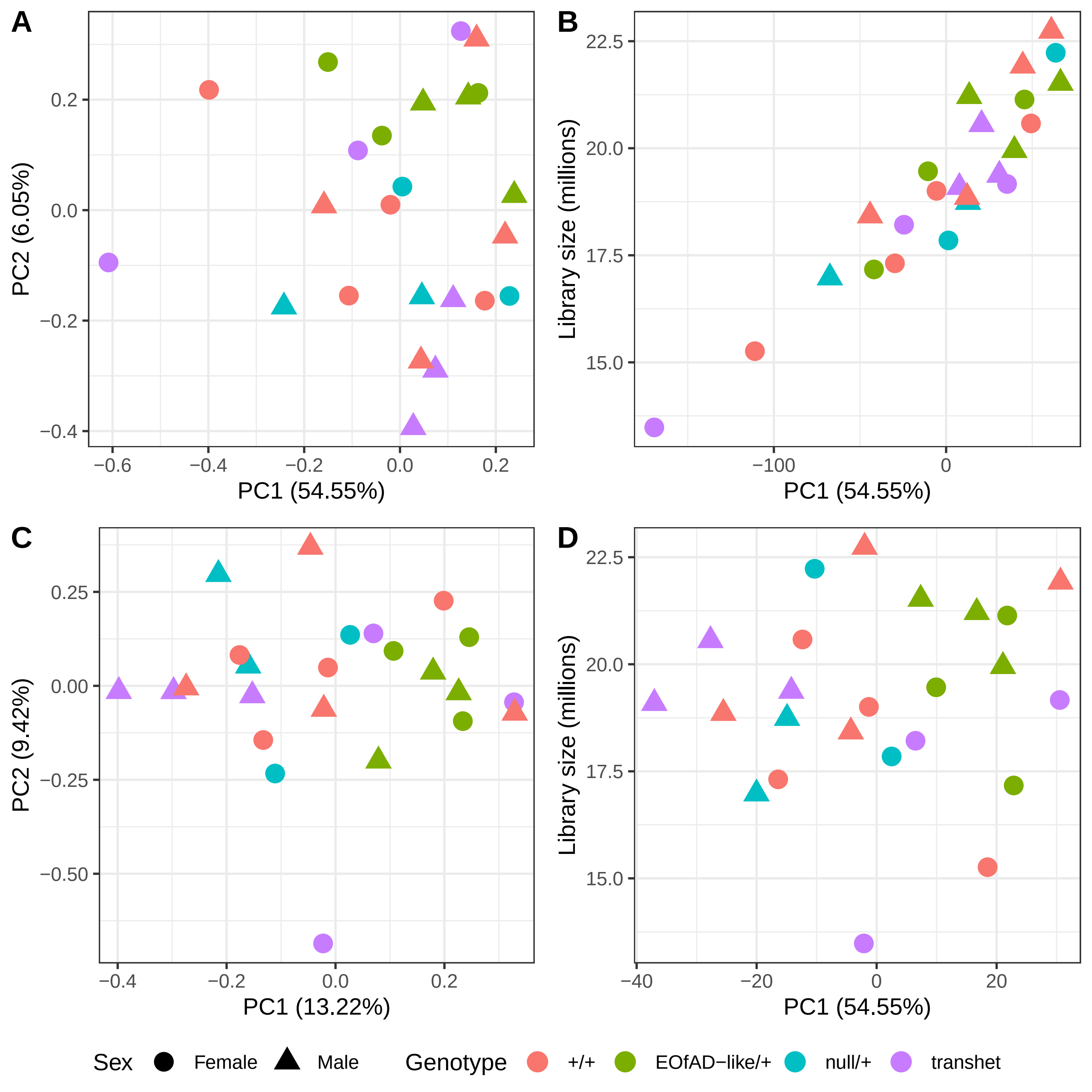

### Additional File 7

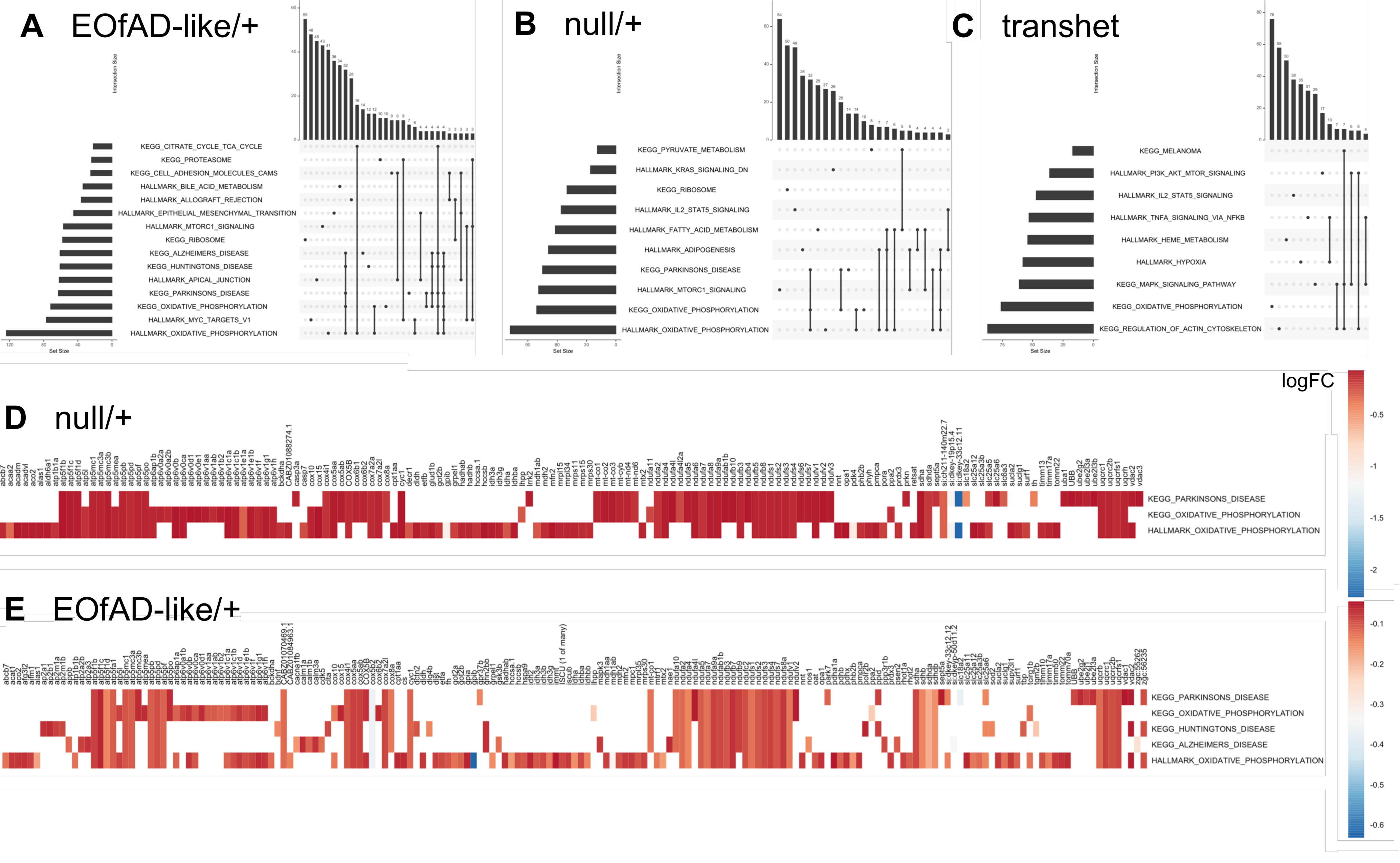

### Additional File 8

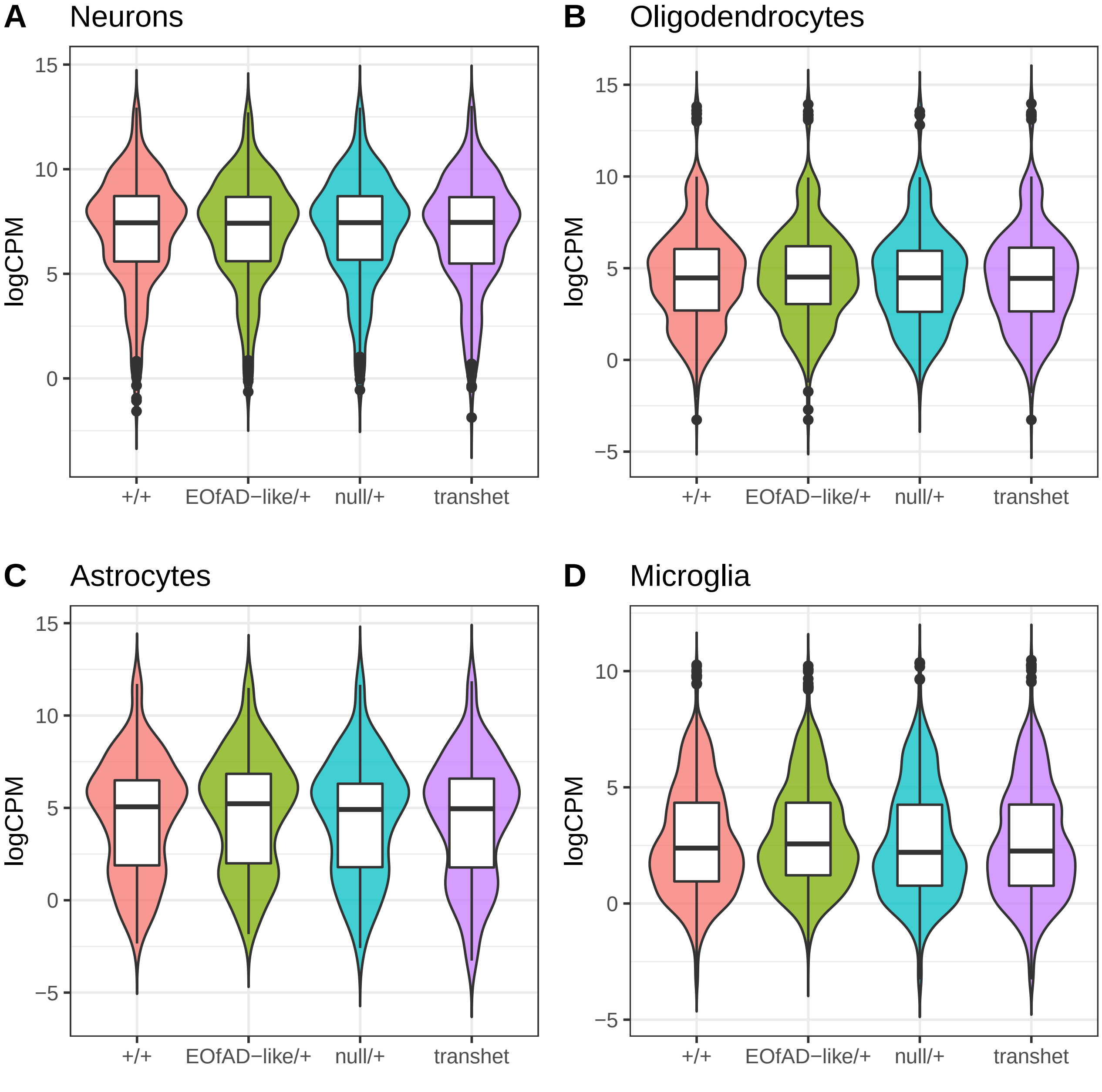

### Additional File 9

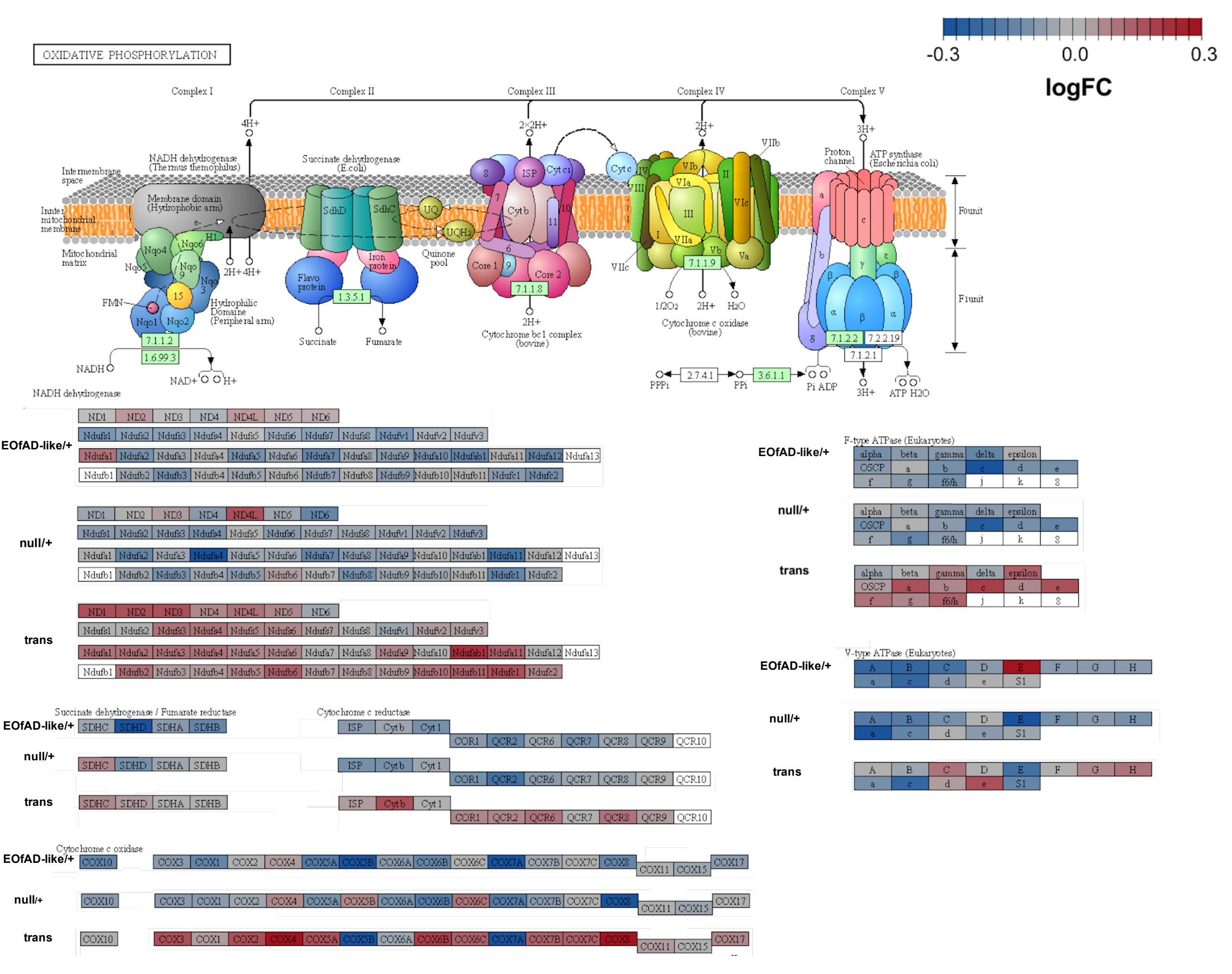

### Additional File 10

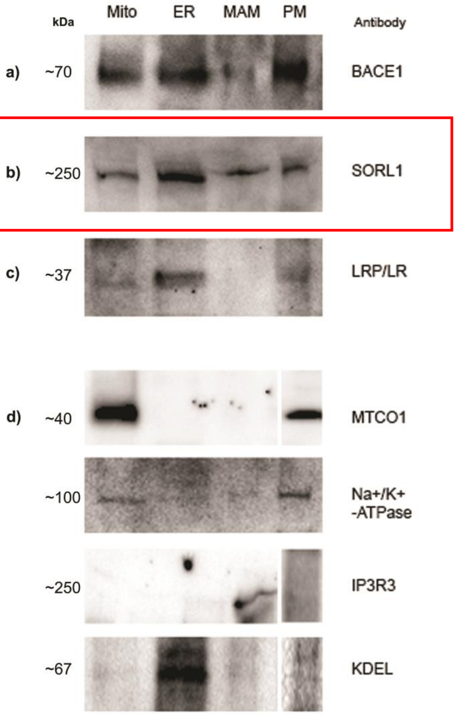
