## Additional File 2 for "Brain transcriptome analysis reveals subtle effects on mitochondrial function and iron homeostasis of mutations in the *SORL1* gene implicated in early onset familial Alzheimer’s disease"

### Additional File 2: Generation of mutant zebrafish lines

A schematic of the generation of mutant lines of zebrafish is found in Figure 1 and detailed below.

### Microinjection into zebrafish embryos

The mRNA solutions encoding the left and right TALENs were diluted to 400 ng/μL in milliQ water, then 2 μLs of each were mixed together to give a final concentration of 200 ng/μL of each TALEN mRNA. 2-3 nL of this solution was injected into Tübingen strain zebrafish embryos at the one cell stage. Injected embryos were incubated at 28°C.

For CRISPR-Cpf1 editing, the crRNA was provided as a lyophilised solid. We re-suspended the crRNA in 1 x tris-EDTA (TE) to give a 100 µM solution. We then prepared a ribonucleoprotein (RNP) solution containing 24 µM crRNA and 20 µM recombinant Cpf1 protein and incubated this at 37°C for 10 minutes as recommended by (Moreno-Mateos *et al.* 2017) . 2-5 nL of the RNP solution was injected into zebrafish embryos at the one cell stage. Injected embryos were incubated at 34 °C until 24 hours post fertilisation (hpf), and then moved to 28 °C.

### Genomic DNA extraction

To extract genomic DNA from embryos, we randomly selected 10 embryos at 24 hpf and placed those in 45 µL of 1 x TE buffer. For tail clips, we anaesthetised the adult fish in Tricaine solution (Westerfield 2007) and clipped off a small piece of the tail. We then added this tail clip to 45 µL of 1 x TE buffer.

To extract the genomic DNA, 5 µL of recombinant Proteinase K solution at 20mg/mL (Roche, Basel, Switzerland) was added to the pooled embryos or to each tail clip and incubated at 55 °C for 3 hours. Proteinase K was inactivated by incubation at 95 °C for 5 minutes and then debris was pelleted by centrifugation with a relative centrifugal force of 16,100 for 3 minutes. The supernatant containing the genomic DNA was then transferred to a clean tube.

### PCR amplification across mutation regions

We amplified across the mutation regions using polymerase chain reactions (PCRs) with GoTaq® DNA Polymerase (Promega, Wisconsin, USA) following the manufacturers protocol. Primers used to these regions are found in Table 1. Cycling conditions consisted of 2 minutes of 95 °C, then 30 cycles of 95 °C for 30 seconds, 60 °C for 30 seconds and 72 °C for 30 seconds, then a final step of 72°C for 5 minutes. PCR products were then separated by electrophoresis through a 1% agarose in TAE buffer gel for 30 minutes at 90 volts. DNA products of interest were cut out using sterile blades and purified from the gel using the Wizard® SV Gel and PCR Clean-Up System (Promega, Wisconsin, USA) following the manufacturer’s protocol. Recovered DNA concentrations were estimated using a Nanodrop spectrophotometer.

| Table 1: Primers used to amplify across mutation regions. | | |
| --- | --- | --- |
| Primer Name | Sequence (5’ to 3’) | Comment |
| *sorl1* exon 2 forward | CGGTGAGAGAAGCAGAAACTAAA | Amplify *sorl1* exon 2 region |
| *sorl1* exon 2 reverse | GCCCATCAGAAGAACCAGAA |  |
| *sorl1* exon 32 forward | CCACATCATCACTGTTCATTTGC | Amplify *sorl1* exon 32 region |
| *sorl1* exon 32 reverse | GCGAAGATGCACAAGGGAA |  |

### T7 endonuclease assay

To determine whether any mutations were present in embryos or tail clips, we used 200 ng of purified DNA amplified from the region spanning the mutagenesis site in a reaction using T7 endonuclease I (T7E1, NEB, Ipswich, USA) following the manufacturer’s protocol.

### Breeding strategy to generate F2 fish

Since the injected, G0 fish are mosaic for any mutations, we pair-mated them with Tübingen fish of approximately the same age to generate F1 families containing individuals either fully heterozygous mutant, or wild-type. To determine whether any mutations were passed on from the G0 fish, we removed a subset of each F1 embryo clutch at 24 hpf for genomic DNA extraction, PCR amplification across the mutation sites and T7EI digestion. Any clutches for which the expected cleavage product sizes after the digestion were seen were allowed to develop into the F1 fish.

We identified mutant fish in F1 families by tail clipping , extraction of genomic DNA, PCR amplification across the mutation sites and then T7EI assays. The PCR products from mutant fish were then cloned into the pGem-T easy vector (Promega, Wisconsin, USA) to isolate the mutant allele, before Sanger sequencing at the Australian Genome Research Facility (AGRF, Adelaide, AUS) after following their recommended protocol for sample preparation. We sequenced in both the forward and reverse directions using M13 primers (M13 forward primer sequence: 5’ GTAAAACGACGGCCAGT 3’. M13 reverse primer sequence 5’ GGAAACAGCTARGACCATG 3’).

Once mutations were identified with predicted premature termination codons close to the mutation site (V1482Afs for exon 32 and R122Pfs for exon 2), we designed allele specific PCR primers to assist with genotyping. The allele specific primers are described in Table 2.

F2 families were generated by outbreeding heterozygous mutant fish with Tübingen fish of approximately the same age.

| Table 2: Allele specific primer sequences | |
| --- | --- |
| Primer | Sequence (5’ - 3’) |
| *sorl1* R122Pfs mutation-specific R | TCTGTGGCGCTAACGCTACTAT |
| *sorl1* R122 WT specific R | CGCTGTCTCTCGCTAACGC |
| *sorl1* R122 common F | CGGTGAGAGAAGCAGAAACTAAA |
| *sorl1* V1482Afs mutation-specific F | AGATCAGTTTCTGTGTGCCTCC |
| *sorl1* V1482 WT specific F | GTGTGTGAAGCCTCCAGCAT |
| *sorl1* V1482 common reverse | GCGAAGATGCACAAGGGAA |

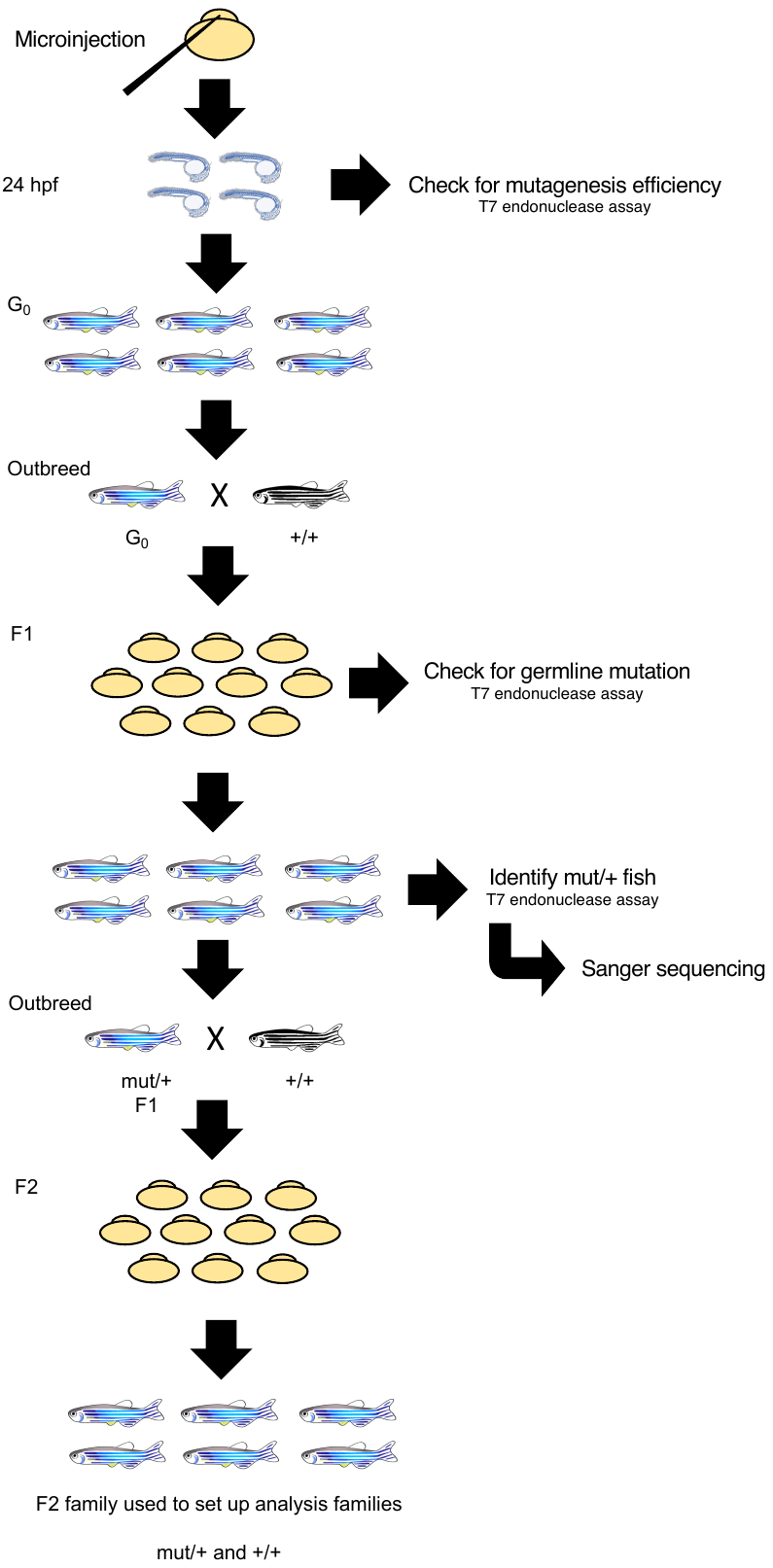

Figure 1: Breeding strategy to generate F2 fish.
