## Additional File 6 for "Brain transcriptome analysis reveals subtle effects on mitochondrial function and iron homeostasis of mutations in the *SORL1* gene implicated in early onset familial Alzheimer’s disease"

**Additional File 2: Gene set enrichment analysis**

This document contains the outputs of the gene set enrichment analyses using fry, camera and fgsea.

### Supplemental Table 1: Top 10 significant KEGG gene sets in each comparison of *sorl1* genotypes with wild type using fry.

| Gene Set | Number of genes | Direction | p | FDR | p mixed | FDR Mixed | Comparison |
| --- | --- | --- | --- | --- | --- | --- | --- |
| KEGG STARCH AND SUCROSE METABOLISM | 29 | Down | 0.01 | 0.91 | 0.29 | 0.84 | null/+ |
| KEGG NICOTINATE AND NICOTINAMIDE METABOLISM | 17 | Down | 0.01 | 0.91 | 0.74 | 0.85 | null/+ |
| KEGG AMINO SUGAR AND NUCLEOTIDE SUGAR METABOLISM | 40 | Down | 0.02 | 0.91 | 0.27 | 0.84 | null/+ |
| KEGG PYRUVATE METABOLISM | 38 | Down | 0.03 | 0.91 | 0.12 | 0.84 | null/+ |
| KEGG PRIMARY IMMUNODEFICIENCY | 11 | Up | 0.04 | 0.91 | 0.83 | 0.90 | null/+ |
| KEGG GLYCOLYSIS GLUCONEOGENESIS | 57 | Down | 0.04 | 0.91 | 0.13 | 0.84 | null/+ |
| KEGG TERPENOID BACKBONE BIOSYNTHESIS | 13 | Down | 0.04 | 0.91 | 0.35 | 0.84 | null/+ |
| KEGG PENTOSE AND GLUCURONATE INTERCONVERSIONS | 18 | Down | 0.05 | 0.91 | 0.31 | 0.84 | null/+ |
| KEGG VIBRIO CHOLERAE INFECTION | 54 | Down | 0.06 | 0.91 | 0.35 | 0.84 | null/+ |
| KEGG GLYCOSPHINGOLIPID BIOSYNTHESIS GANGLIO SERIES | 16 | Up | 0.06 | 0.91 | 0.00 | 0.35 | null/+ |
| KEGG MELANOMA | 62 | Down | 0.00 | 0.19 | 0.14 | 0.44 | trans |
| KEGG RNA POLYMERASE | 25 | Up | 0.00 | 0.28 | 0.03 | 0.44 | trans |
| KEGG N GLYCAN BIOSYNTHESIS | 46 | Up | 0.02 | 0.85 | 0.04 | 0.44 | trans |
| KEGG GLIOMA | 67 | Down | 0.02 | 0.85 | 0.08 | 0.44 | trans |
| KEGG SPLICEOSOME | 130 | Up | 0.03 | 0.85 | 0.07 | 0.44 | trans |
| KEGG PYRIMIDINE METABOLISM | 80 | Up | 0.04 | 0.85 | 0.04 | 0.44 | trans |
| KEGG THYROID CANCER | 32 | Down | 0.06 | 0.85 | 0.14 | 0.44 | trans |
| KEGG B CELL RECEPTOR SIGNALING PATHWAY | 68 | Down | 0.06 | 0.85 | 0.20 | 0.44 | trans |
| KEGG PRION DISEASES | 25 | Down | 0.08 | 0.85 | 0.09 | 0.44 | trans |
| KEGG BASAL TRANSCRIPTION FACTORS | 29 | Up | 0.08 | 0.85 | 0.61 | 0.70 | trans |
| KEGG NATURAL KILLER CELL MEDIATED CYTOTOXICITY | 83 | Up | 0.00 | 0.14 | 0.29 | 0.57 | EOfAD-like/+ |
| KEGG AMYOTROPHIC LATERAL SCLEROSIS ALS | 54 | Up | 0.01 | 0.40 | 0.39 | 0.58 | EOfAD-like/+ |
| KEGG CELL ADHESION MOLECULES CAMS | 91 | Up | 0.01 | 0.40 | 0.06 | 0.52 | EOfAD-like/+ |
| KEGG VEGF SIGNALING PATHWAY | 68 | Up | 0.01 | 0.50 | 0.14 | 0.52 | EOfAD-like/+ |
| KEGG NON SMALL CELL LUNG CANCER | 53 | Up | 0.02 | 0.53 | 0.18 | 0.52 | EOfAD-like/+ |
| KEGG CITRATE CYCLE TCA CYCLE | 34 | Down | 0.02 | 0.53 | 0.05 | 0.52 | EOfAD-like/+ |
| KEGG ADHERENS JUNCTION | 92 | Up | 0.02 | 0.53 | 0.04 | 0.52 | EOfAD-like/+ |
| KEGG FC EPSILON RI SIGNALING PATHWAY | 76 | Up | 0.02 | 0.53 | 0.22 | 0.53 | EOfAD-like/+ |
| KEGG NOTCH SIGNALING PATHWAY | 54 | Up | 0.03 | 0.53 | 0.14 | 0.52 | EOfAD-like/+ |
| KEGG FC GAMMA R MEDIATED PHAGOCYTOSIS | 98 | Up | 0.03 | 0.53 | 0.05 | 0.52 | EOfAD-like/+ |

### Supplemental Table 2: Top 10 significant HALLMARK gene sets in each comparison of *sorl1* genotypes with wild type using fry.

| Gene Set | Number of genes | Direction | p | FDR | p mixed | FDR Mixed | Comparison |
| --- | --- | --- | --- | --- | --- | --- | --- |
| HALLMARK MTORC1 SIGNALING | 210 | Down | 0.07 | 0.93 | 0.28 | 0.81 | null/+ |
| HALLMARK KRAS SIGNALING DN | 134 | Up | 0.09 | 0.93 | 0.39 | 0.81 | null/+ |
| HALLMARK ANDROGEN RESPONSE | 98 | Down | 0.09 | 0.93 | 0.32 | 0.81 | null/+ |
| HALLMARK PI3K AKT MTOR SIGNALING | 104 | Down | 0.15 | 0.93 | 0.33 | 0.81 | null/+ |
| HALLMARK OXIDATIVE PHOSPHORYLATION | 225 | Down | 0.16 | 0.93 | 0.52 | 0.81 | null/+ |
| HALLMARK MYC TARGETS V1 | 219 | Down | 0.17 | 0.93 | 0.46 | 0.81 | null/+ |
| HALLMARK GLYCOLYSIS | 199 | Down | 0.21 | 0.93 | 0.34 | 0.81 | null/+ |
| HALLMARK SPERMATOGENESIS | 109 | Down | 0.22 | 0.93 | 0.31 | 0.81 | null/+ |
| HALLMARK CHOLESTEROL HOMEOSTASIS | 79 | Down | 0.23 | 0.93 | 0.35 | 0.81 | null/+ |
| HALLMARK FATTY ACID METABOLISM | 158 | Down | 0.24 | 0.93 | 0.20 | 0.81 | null/+ |
| HALLMARK PI3K AKT MTOR SIGNALING | 104 | Down | 0.04 | 0.92 | 0.07 | 0.38 | trans |
| HALLMARK DNA REPAIR | 148 | Up | 0.05 | 0.92 | 0.19 | 0.39 | trans |
| HALLMARK HEME METABOLISM | 179 | Down | 0.06 | 0.92 | 0.11 | 0.38 | trans |
| HALLMARK IL2 STAT5 SIGNALING | 170 | Down | 0.13 | 0.92 | 0.02 | 0.38 | trans |
| HALLMARK REACTIVE OXYGEN SPECIES PATHWAY | 44 | Up | 0.17 | 0.92 | 0.24 | 0.43 | trans |
| HALLMARK APICAL SURFACE | 37 | Down | 0.19 | 0.92 | 0.62 | 0.64 | trans |
| HALLMARK TNFA SIGNALING VIA NFKB | 164 | Down | 0.21 | 0.92 | 0.05 | 0.38 | trans |
| HALLMARK UNFOLDED PROTEIN RESPONSE | 119 | Up | 0.22 | 0.92 | 0.16 | 0.38 | trans |
| HALLMARK MYC TARGETS V1 | 219 | Up | 0.28 | 0.92 | 0.16 | 0.38 | trans |
| HALLMARK OXIDATIVE PHOSPHORYLATION | 225 | Up | 0.29 | 0.92 | 0.31 | 0.51 | trans |
| HALLMARK INFLAMMATORY RESPONSE | 107 | Up | 0.03 | 0.63 | 0.17 | 0.43 | EOfAD-like/+ |
| HALLMARK KRAS SIGNALING UP | 162 | Up | 0.04 | 0.63 | 0.07 | 0.41 | EOfAD-like/+ |
| HALLMARK APICAL JUNCTION | 191 | Up | 0.05 | 0.63 | 0.11 | 0.41 | EOfAD-like/+ |
| HALLMARK KRAS SIGNALING DN | 134 | Up | 0.06 | 0.63 | 0.62 | 0.71 | EOfAD-like/+ |
| HALLMARK ALLOGRAFT REJECTION | 112 | Up | 0.08 | 0.63 | 0.11 | 0.41 | EOfAD-like/+ |
| HALLMARK HEDGEHOG SIGNALING | 44 | Up | 0.10 | 0.63 | 0.34 | 0.54 | EOfAD-like/+ |
| HALLMARK BILE ACID METABOLISM | 96 | Up | 0.11 | 0.63 | 0.09 | 0.41 | EOfAD-like/+ |
| HALLMARK UV RESPONSE DN | 154 | Up | 0.12 | 0.63 | 0.22 | 0.45 | EOfAD-like/+ |
| HALLMARK INTERFERON ALPHA RESPONSE | 56 | Up | 0.14 | 0.63 | 0.51 | 0.67 | EOfAD-like/+ |
| HALLMARK OXIDATIVE PHOSPHORYLATION | 225 | Down | 0.16 | 0.63 | 0.16 | 0.43 | EOfAD-like/+ |

### Supplemental Table 3: Top 10 significant KEGG gene sets in each comparison of *sorl1* genotypes with wild type using camera.

| Gene set | Number of genes | Correlation | Direction | p | FDR | Comparison |
| --- | --- | --- | --- | --- | --- | --- |
| KEGG STARCH AND SUCROSE METABOLISM | 29 | 0.02 | Down | 0.00 | 0.65 | null/+ |
| KEGG AMINO SUGAR AND NUCLEOTIDE SUGAR METABOLISM | 40 | 0.01 | Down | 0.01 | 0.65 | null/+ |
| KEGG NICOTINATE AND NICOTINAMIDE METABOLISM | 17 | -0.02 | Down | 0.02 | 0.88 | null/+ |
| KEGG GLYCOLYSIS GLUCONEOGENESIS | 57 | 0.03 | Down | 0.03 | 0.88 | null/+ |
| KEGG PENTOSE AND GLUCURONATE INTERCONVERSIONS | 18 | 0.03 | Down | 0.03 | 0.88 | null/+ |
| KEGG PYRUVATE METABOLISM | 38 | 0.10 | Down | 0.03 | 0.88 | null/+ |
| KEGG TERPENOID BACKBONE BIOSYNTHESIS | 13 | 0.10 | Down | 0.04 | 0.88 | null/+ |
| KEGG PRIMARY BILE ACID BIOSYNTHESIS | 18 | 0.07 | Down | 0.05 | 0.90 | null/+ |
| KEGG VIBRIO CHOLERAE INFECTION | 54 | 0.01 | Down | 0.05 | 0.90 | null/+ |
| KEGG PRIMARY IMMUNODEFICIENCY | 11 | -0.01 | Up | 0.07 | 0.90 | null/+ |
| KEGG MELANOMA | 62 | 0.00 | Down | 0.00 | 0.22 | trans |
| KEGG RNA POLYMERASE | 25 | 0.04 | Up | 0.01 | 0.80 | trans |
| KEGG GLIOMA | 67 | 0.01 | Down | 0.02 | 0.85 | trans |
| KEGG B CELL RECEPTOR SIGNALING PATHWAY | 68 | 0.01 | Down | 0.03 | 0.85 | trans |
| KEGG THYROID CANCER | 32 | 0.04 | Down | 0.06 | 0.85 | trans |
| KEGG N GLYCAN BIOSYNTHESIS | 46 | 0.00 | Up | 0.07 | 0.85 | trans |
| KEGG ADIPOCYTOKINE SIGNALING PATHWAY | 66 | 0.00 | Down | 0.07 | 0.85 | trans |
| KEGG PYRIMIDINE METABOLISM | 80 | 0.03 | Up | 0.07 | 0.85 | trans |
| KEGG PRION DISEASES | 25 | 0.01 | Down | 0.07 | 0.85 | trans |
| KEGG SPLICEOSOME | 130 | 0.05 | Up | 0.08 | 0.85 | trans |
| KEGG NATURAL KILLER CELL MEDIATED CYTOTOXICITY | 83 | 0.00 | Up | 0.02 | 0.89 | EOfAD-like/+ |
| KEGG CITRATE CYCLE TCA CYCLE | 34 | 0.10 | Down | 0.02 | 0.89 | EOfAD-like/+ |
| KEGG AMINO SUGAR AND NUCLEOTIDE SUGAR METABOLISM | 40 | 0.01 | Down | 0.02 | 0.89 | EOfAD-like/+ |
| KEGG GLYCOLYSIS GLUCONEOGENESIS | 57 | 0.03 | Down | 0.03 | 0.89 | EOfAD-like/+ |
| KEGG VIBRIO CHOLERAE INFECTION | 54 | 0.01 | Down | 0.04 | 0.89 | EOfAD-like/+ |
| KEGG CELL ADHESION MOLECULES CAMS | 91 | 0.02 | Up | 0.04 | 0.89 | EOfAD-like/+ |
| KEGG FRUCTOSE AND MANNOSE METABOLISM | 41 | 0.01 | Down | 0.06 | 0.89 | EOfAD-like/+ |
| KEGG VALINE LEUCINE AND ISOLEUCINE BIOSYNTHESIS | 12 | 0.07 | Down | 0.07 | 0.89 | EOfAD-like/+ |
| KEGG GLYOXYLATE AND DICARBOXYLATE METABOLISM | 15 | 0.05 | Down | 0.07 | 0.89 | EOfAD-like/+ |
| KEGG VEGF SIGNALING PATHWAY | 68 | 0.02 | Up | 0.07 | 0.89 | EOfAD-like/+ |

### Supplemental Table 4: Top 10 significant HALLMARK gene sets in each comparison of *sorl1* genotypes with wild type using camera.

| Gene set | Number of genes | Correlation | Direction | p | FDR | Comparison |
| --- | --- | --- | --- | --- | --- | --- |
| HALLMARK MTORC1 SIGNALING | 210 | 0.05 | Down | 0.04 | 0.73 | null/+ |
| HALLMARK ANDROGEN RESPONSE | 98 | 0.02 | Down | 0.05 | 0.73 | null/+ |
| HALLMARK PI3K AKT MTOR SIGNALING | 104 | 0.00 | Down | 0.06 | 0.73 | null/+ |
| HALLMARK GLYCOLYSIS | 199 | 0.02 | Down | 0.11 | 0.73 | null/+ |
| HALLMARK MYC TARGETS V1 | 219 | 0.05 | Down | 0.13 | 0.73 | null/+ |
| HALLMARK OXIDATIVE PHOSPHORYLATION | 225 | 0.14 | Down | 0.14 | 0.73 | null/+ |
| HALLMARK KRAS SIGNALING DN | 134 | 0.01 | Up | 0.14 | 0.73 | null/+ |
| HALLMARK CHOLESTEROL HOMEOSTASIS | 79 | 0.04 | Down | 0.15 | 0.73 | null/+ |
| HALLMARK SPERMATOGENESIS | 109 | 0.01 | Down | 0.16 | 0.73 | null/+ |
| HALLMARK IL2 STAT5 SIGNALING | 170 | 0.03 | Down | 0.17 | 0.73 | null/+ |
| HALLMARK PI3K AKT MTOR SIGNALING | 104 | 0.00 | Down | 0.02 | 0.83 | trans |
| HALLMARK HEME METABOLISM | 179 | 0.01 | Down | 0.04 | 0.83 | trans |
| HALLMARK IL2 STAT5 SIGNALING | 170 | 0.03 | Down | 0.09 | 0.83 | trans |
| HALLMARK DNA REPAIR | 148 | 0.03 | Up | 0.12 | 0.83 | trans |
| HALLMARK APICAL SURFACE | 37 | 0.02 | Down | 0.14 | 0.83 | trans |
| HALLMARK TNFA SIGNALING VIA NFKB | 164 | 0.04 | Down | 0.16 | 0.83 | trans |
| HALLMARK KRAS SIGNALING UP | 162 | 0.01 | Down | 0.18 | 0.83 | trans |
| HALLMARK REACTIVE OXYGEN SPECIES PATHWAY | 44 | 0.08 | Up | 0.25 | 0.83 | trans |
| HALLMARK SPERMATOGENESIS | 109 | 0.01 | Down | 0.27 | 0.83 | trans |
| HALLMARK IL6 JAK STAT3 SIGNALING | 43 | 0.04 | Down | 0.30 | 0.83 | trans |
| HALLMARK SPERMATOGENESIS | 109 | 0.01 | Down | 0.05 | 0.88 | EOfAD-like/+ |
| HALLMARK OXIDATIVE PHOSPHORYLATION | 225 | 0.14 | Down | 0.09 | 0.88 | EOfAD-like/+ |
| HALLMARK PI3K AKT MTOR SIGNALING | 104 | 0.00 | Down | 0.09 | 0.88 | EOfAD-like/+ |
| HALLMARK UNFOLDED PROTEIN RESPONSE | 119 | 0.03 | Down | 0.13 | 0.88 | EOfAD-like/+ |
| HALLMARK MYC TARGETS V1 | 219 | 0.05 | Down | 0.17 | 0.88 | EOfAD-like/+ |
| HALLMARK INFLAMMATORY RESPONSE | 107 | 0.02 | Up | 0.17 | 0.88 | EOfAD-like/+ |
| HALLMARK APICAL JUNCTION | 191 | 0.03 | Up | 0.19 | 0.88 | EOfAD-like/+ |
| HALLMARK UV RESPONSE UP | 170 | 0.02 | Down | 0.19 | 0.88 | EOfAD-like/+ |
| HALLMARK GLYCOLYSIS | 199 | 0.02 | Down | 0.20 | 0.88 | EOfAD-like/+ |
| HALLMARK MYC TARGETS V2 | 60 | 0.05 | Down | 0.21 | 0.88 | EOfAD-like/+ |

### Supplemental Table 5: Significant KEGG gene sets in each comparison of *sorl1* genotypes with wild type using fgsea.

| Gene set | p | p_bonferroni_ | NES | Number of genes | Comparison |
| --- | --- | --- | --- | --- | --- |
| KEGG RIBOSOME | 0.00 | 0.00 | -2.29 | 71 | null/+ |
| KEGG OXIDATIVE PHOSPHORYLATION | 0.00 | 0.00 | -2.23 | 120 | null/+ |
| KEGG PARKINSONS DISEASE | 0.00 | 0.00 | -2.07 | 123 | null/+ |
| KEGG PYRUVATE METABOLISM | 0.00 | 0.01 | -2.26 | 38 | null/+ |
| KEGG OXIDATIVE PHOSPHORYLATION | 0.00 | 0.00 | 2.05 | 120 | trans |
| KEGG OXIDATIVE PHOSPHORYLATION | 0.00 | 0.00 | -2.45 | 120 | EOfAD-like/+ |
| KEGG PARKINSONS DISEASE | 0.00 | 0.00 | -2.07 | 123 | EOfAD-like/+ |
| KEGG RIBOSOME | 0.00 | 0.00 | 2.02 | 71 | EOfAD-like/+ |
| KEGG CITRATE CYCLE TCA CYCLE | 0.00 | 0.00 | -2.27 | 34 | EOfAD-like/+ |
| KEGG PROTEASOME | 0.00 | 0.00 | -2.20 | 47 | EOfAD-like/+ |
| KEGG ALZHEIMERS DISEASE | 0.00 | 0.01 | -1.80 | 151 | EOfAD-like/+ |
| KEGG CELL ADHESION MOLECULES CAMS | 0.00 | 0.01 | 1.85 | 91 | EOfAD-like/+ |

### Supplemental Table 6: Significant HALLMARK gene sets in each comparison of *sorl1* genotypes with wild type using fgsea

| Gene set | p | p_bonferroni_ | NES | Number of genes | Comparison |
| --- | --- | --- | --- | --- | --- |
| HALLMARK MTORC1 SIGNALING | 0.00 | 0.00 | -2.04 | 210 | null/+ |
| HALLMARK OXIDATIVE PHOSPHORYLATION | 0.00 | 0.00 | -2.22 | 225 | null/+ |
| HALLMARK IL2 STAT5 SIGNALING | 0.00 | 0.00 | -1.92 | 178 | null/+ |
| HALLMARK FATTY ACID METABOLISM | 0.00 | 0.00 | -1.87 | 159 | null/+ |
| HALLMARK ADIPOGENESIS | 0.00 | 0.00 | -1.77 | 204 | null/+ |
| HALLMARK KRAS SIGNALING DN | 0.00 | 0.03 | 1.67 | 135 | null/+ |
| HALLMARK HEME METABOLISM | 0.00 | 0.00 | -1.84 | 191 | trans |
| HALLMARK TNFA SIGNALING VIA NFKB | 0.00 | 0.00 | -1.79 | 167 | trans |
| HALLMARK IL2 STAT5 SIGNALING | 0.00 | 0.00 | -1.73 | 178 | trans |
| HALLMARK MYC TARGETS V1 | 0.00 | 0.00 | -1.81 | 219 | EOfAD-like/+ |
| HALLMARK OXIDATIVE PHOSPHORYLATION | 0.00 | 0.00 | -2.37 | 225 | EOfAD-like/+ |
| HALLMARK ALLOGRAFT REJECTION | 0.00 | 0.00 | 1.89 | 117 | EOfAD-like/+ |
| HALLMARK BILE ACID METABOLISM | 0.00 | 0.01 | 1.74 | 99 | EOfAD-like/+ |
| HALLMARK EPITHELIAL MESENCHYMAL TRANSITION | 0.00 | 0.03 | 1.57 | 190 | EOfAD-like/+ |
| HALLMARK MTORC1 SIGNALING | 0.00 | 0.03 | -1.55 | 210 | EOfAD-like/+ |
